# Type 2 diabetes amplifies environmental Pb toxicity and intergenerational risk

**DOI:** 10.64898/2026.02.21.707175

**Authors:** Chenyi Bo Zhang, Xiaoping Xia, Chen Zhang, Chenyu Pei, Zhixiang Guo, Guangnan Li, Liang Xu Ma, Huali Luo, Jiaoyan Ma, Jingwen Luo, Hongxiao Cui, Banghua He, Qin Li, Yukai Jason Zhong, Haifeng Zhang, Xianguang Yang, Xin Zhiguo Li

**Author notes:** These authors contributed equally to this work (Chenyi Bo Zhang, Xiaoping Xia, Chen Zhang).

## Abstract

Type 2 diabetes (T2D) and environmental lead (Pb) exposure intersect as prevalent global health challenges. Prior studies report elevated blood Pb in T2D, but the underlying mechanism and directionality remain unresolved. Here we show that T2D fundamentally alters Pb toxicokinetics, converting otherwise modest exposure into progressive organ injury and heritable risk. In a human cohort, elevated blood Pb in T2D specifically associated with impaired renal function rather than glycaemic control or adiposity. In db/db mice, diabetes increased oral Pb absorption through delayed gastrointestinal transit and epithelial barrier dysfunction, leading to enhanced systemic retention. With sustained exposure, renal Pb accumulation induced tubular stress that reduced effective clearance, creating a feed-forward loop in which diabetes promotes Pb retention and retained Pb accelerates kidney injury. Chronic Pb exposure exacerbated diabetic kidney injury without inducing hyperglycaemia, and short-course chelation attenuated renal stress markers. Beyond somatic toxicity, combined paternal T2D and Pb exposure programmed renal and neurobehavioral phenotypes across generations via sperm RNA. These findings redefine Pb from a passive environmental exposure to a context-dependent driver of disease, identify T2D as a vulnerability state that reshapes toxicant handling, and establish a mechanistic link between metabolic disease, environmental exposure and intergenerational risk. This work highlights the need for vulnerability-informed clinical management and environmental regulation in populations burdened by T2D.

## Introduction

Type 2 diabetes mellitus (T2D) affects more than half billion people worldwide and is a leading cause of kidney failure, cardiovascular morbidity and premature mortality. Despite advances in glycaemic control and renoprotective therapy, substantial heterogeneity in disease progression remains unexplained. Clinical trajectories often diverge markedly among individuals with comparable metabolic control, suggesting that contextual or environmental modifiers contribute to disease severity beyond hyperglycaemia alone^1^.

Lead (Pb) is a pervasive environmental toxicant with no known physiological role and well-documented neurotoxic, nephrotoxic and reproductive effects. Although regulatory measures have reduced population-level exposure, low-level Pb remains detectable in food, water and consumer products and disproportionately affects communities with low socioeconomic status^2^. Epidemiological surveys estimate that up to **800 million** children worldwide have blood-lead concentrations ≥5 µg dl⁻¹, and more than half of the U.S. population were exposed to harmful lead levels during early childhood^3^. Experimental studies further show that maternal Pb exposure can induce persistent epigenetic and behavioral alterations across generations^4,5^, raising the possibility that environmental Pb may exert long-term biological effects beyond direct toxicity.

Several epidemiological studies have reported elevated blood Pb levels in individuals with T2D^6–9^. A longitudinal analysis suggests that higher body Pb burden predicts accelerated decline in estimated glomerular filtration rate (eGFR) among patients with T2D, and an early interventional study reported that chelation therapy with calcium disodium EDTA slowed progression of diabetic nephropathy in selected cohorts^10,11^. Together, these findings imply that Pb burden may modify the trajectory of diabetic kidney disease. However, the mechanistic basis of the Pb–T2D association and whether Pb interact with T2D to shape organ vulnerability and inherited disease risk remains unclear.

These observations raise three non-exclusive possibilities: (i) individuals with T2D experience greater environmental Pb exposure; (ii) Pb contributes causally to diabetes development; or (iii) the diabetic state alters Pb absorption/clearance. To disentangle these scenarios, we combined analyses of human cohorts with diabetic mouse models and multigenerational experiments, revealing that diabetes markedly increases oral lead uptake, accelerates renal and testicular injury, and increases neurobehavioral and renal susceptibility across generations via sperm RNA. Together, our findings reveal a vicious cycle in which T2D amplifies lead toxicity and underscore the need for disease-specific exposure guidelines and targeted chelation strategies.

## Results

### Elevated blood Pb burden in T2D

We quantified whole-blood Pb concentrations by inductively coupled plasma–mass spectrometry (ICP–MS) in 118 individuals with T2D and 61 age- and sex-matched non-diabetic controls (Extended Data Fig. 1a). Blood Pb concentrations were significantly higher in individuals with T2D and displayed greater inter-individual variability compared with controls (Fig. 1a). Seventeen percent of individuals with T2D exceeded the U.S. Centers for Disease Control and Prevention (CDC) blood Pb reference value of 3.5 μg dL⁻¹, whereas none of the non-diabetic controls did (Fig. 1a). This reference value is used for public health surveillance and does not represent a clinical toxicity threshold. Elevated Pb concentrations were observed in both male and female participants with T2D (Fig. 1a).

**Figure 1.**
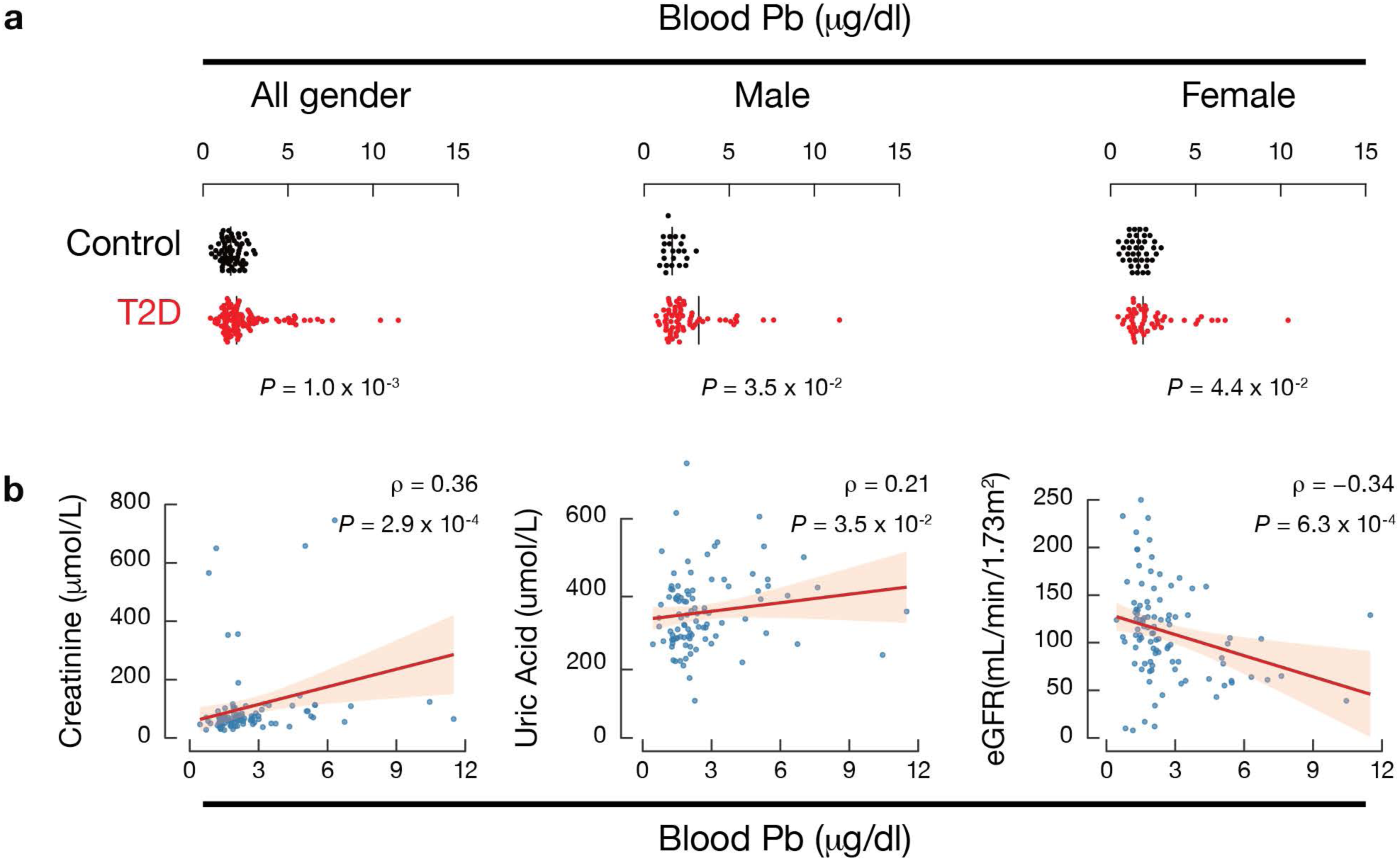
Elevated blood Pb burden in type 2 diabetes. **a**, Whole-blood Pb concentrations in non-diabetic controls and individuals with T2D, shown for all participants and stratified by sex. Each point represents one individual. Data are mean ± s.e.m. Two-tailed Mann–Whitney U test. **b**, Spearman correlation analysis between blood Pb and clinical parameters (creatinine, uric acid, eGFR) in the T2D cohort (n = 98 with complete renal function data). Heat map displays Spearman’s ρ values. *P* values indicate significance of correlation.

To assess clinical relevance while minimizing confounding across groups, we performed correlation analyses separately within the T2D and non-diabetic control cohorts. Among individuals with T2D, blood Pb concentrations were positively correlated with serum creatinine and uric acid, and inversely correlated with estimated glomerular filtration rate (eGFR) (Fig. 1b). These associations were not detected in non-diabetic controls (Extended Data Fig. 1b). No significant correlations were observed between blood Pb and body-mass index (*P* = 0.91) or liver-related biochemical parameters, including serum albumin, triglycerides and total cholesterol (*P* ≥ 0.10; Extended Data Fig. 1b). Together, these findings indicate that elevated Pb burden in T2D co-varies with impaired renal function rather than with adiposity or hepatic dysfunction.

To assess whether elevated Pb is broadly associated with dysregulated glucose metabolism or preferentially observed in adult-onset T2D, we examined additional diabetes subgroups, including type 1 diabetes, juvenile-onset T2D, and gestational diabetes mellitus. In these groups, blood Pb concentrations were not significantly different from age-matched non-diabetic controls (Extended Data Fig. 1c). These findings suggest that elevated Pb burden is not a uniform feature across all forms of diabetes and may preferentially associate with the metabolic context characteristic of adult-onset T2D.

Multi-element ICP–MS profiling of a subset of samples (15 of 27 detectable elements) revealed that Pb showed the most pronounced and statistically significant elevation in T2D (*P* = 2.0 × 10⁻⁴), whereas other metals exhibited modest or no differences (Extended Data Fig. 1d; Table 1). These data support a selective alteration of Pb burden in T2D rather than generalized metal dysregulation.

### Diabetes increases systemic Pb accumulation

To investigate the mechanistic basis of elevated blood Pb in T2D, we employed the db/db mouse model, which carries a loss-of-function mutation in the leptin receptor and develops obesity, hyperglycaemia and early diabetic kidney changes^12^, together with age-matched non-diabetic littermate controls from the same genetic background (m/m). Male mice received 100 ppm lead (II) acetate in drinking water beginning at 8 weeks of age and continuing for 8 weeks (Fig. 2a). This exposure concentration has been widely used in experimental Pb toxicology studies^5,13–16^. In non-diabetic controls, this regimen produced steady-state blood Pb concentrations of 7.0 ± 1.0 μg dL⁻¹ (Fig. 2b, left), levels comparable to those reported in environmentally exposed populations residing in contaminated housing or legacy industrial settings^17^. In db/db mice, blood Pb concentrations reached 30 ± 6 μg dL⁻¹ (Fig. 2b), remaining below the Occupational Safety and Health Administration (OSHA) medical removal level for adult occupational exposure (50 μg dL⁻¹).

**Figure 2.**
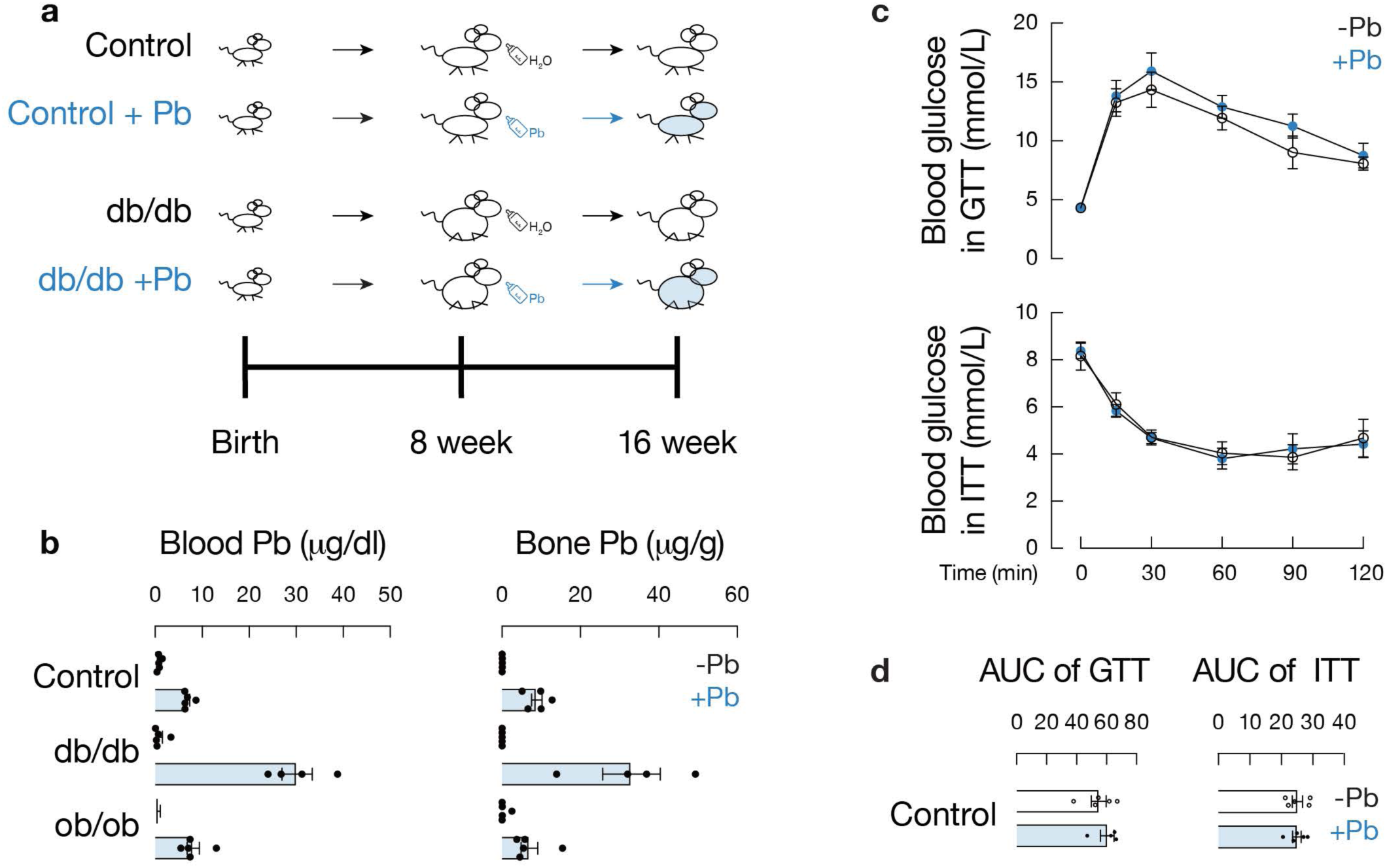
Diabetes increases systemic Pb accumulation without altering glycaemic control. **a**, Experimental design. Male non-diabetic controls (m/m) and db/db mice received 0 or lead(II) acetate (100 ppm) in drinking water from 8 to 16 weeks of age. **b**, Whole-blood Pb concentration and femoral bone Pb content at week 16 in m/m, m/m(Pb), db/db, db/db(Pb), and ob/ob mice (n = 4-5 per group). **c**, Intraperitoneal glucose tolerance test (IPGTT; 16-h fast) and insulin tolerance test (ITT; 4-h fast) at week 16. **d**, Area under the curve (AUC) analysis for IPGTT and ITT. Data are mean ± s.e.m. Two-way ANOVA with post hoc correction unless otherwise indicated. n.s., not significant.

Paternal exposure was initiated in adulthood (8 weeks of age) and maintained for 8 weeks to model environmentally acquired risk while avoiding confounding developmental effects. Restricting exposure to adulthood isolates germline transmission from prenatal and maternal influences and follows established paradigms demonstrating that adult exposure induces heritable epigenetic changes^18–21^. The 8-week exposure was selected to model sustained environmental Pb exposure, allowing equilibration of Pb in blood and bone^5,13,14^, while spanning multiple cycles of spermatogenesis to enable transgenerational analysis.

Following exposure, db/db mice accumulated significantly higher blood Pb levels than non-diabetic controls (*P* = 8.6 × 10⁻^5^) (Fig. 2b). Serial monitoring demonstrated that blood Pb in db/db mice was already elevated by day 1 (*P* = 0.017), reached a steady state by week 2, and remained stable thereafter (Extended Data Fig. 2a). Bone Pb concentrations, representing the major long-term reservoir^22^, were similarly increased in db/db mice (33 ± 15 μg g⁻¹ versus 9 ± 3 μg g⁻¹; *P* = 8.2 × 10⁻^3^, Fig. 2b), indicating that the diabetic state shifts systemic Pb equilibrium to a higher set point.

To determine whether increased Pb burden was attributable to obesity or increased water intake, we examined obese but non-diabetic ob/ob mice^23,24^. Despite greater body weight (61 ± 2 g versus 44 ± 5 g in db/db, Extended Data Fig. 2b), ob/ob mice did not exhibit elevated blood or bone Pb compared with non-diabetic controls (*P* = 0.41 and 0.48, respectively, Fig. 2b). These findings indicate that obesity alone is insufficient to drive enhanced Pb accumulation and instead implicate the diabetic metabolic state as the critical determinant. This interpretation is consistent with the absence of correlation between blood Pb and body-mass index in individuals with T2D (Extended Data Fig. 1b).

To address directionality, we next tested whether chronic Pb exposure worsened glycaemic control. Pb exposure did not significantly alter body weight, fasting glucose, glucose tolerance, or insulin sensitivity in either db/db or non-diabetic control mice (Fig. 2c, d; Extended Data Fig. 2c; *P* ≥ 0.41). These data support a model in which diabetes enhances systemic Pb retention rather than Pb serving as a primary driver of hyperglycaemia. Consistent with this interpretation, blood Pb did not correlate with glycated hemoglobin (HbA1c) or fasting plasma glucose in the human T2D cohort (Extended Data Fig. 1b).

### Diabetes enhances oral Pb absorption

We next examined whether diabetes alters Pb toxicokinetics by affecting intestinal uptake or systemic clearance. To dissociate absorption from elimination, we compared blood Pb kinetics following intravenous versus oral administration (Fig. 3a, Extended Data Fig. 3a). After intravenous injection, which bypasses intestinal uptake, blood Pb clearance was indistinguishable between db/db and non-diabetic control mice, indicating preserved systemic elimination capacity. In contrast, oral gavage with a fixed Pb dose normalized to body weight produced a distinct kinetic profile. Db/db mice exhibited higher blood Pb concentrations at 2 hours post-gavage compared with controls. Notably, non-diabetic controls displayed an earlier peak at 30 minutes, whereas db/db mice exhibited delayed yet sustained elevation, indicating altered intestinal absorption dynamics rather than impaired systemic clearance.

**Figure 3.**
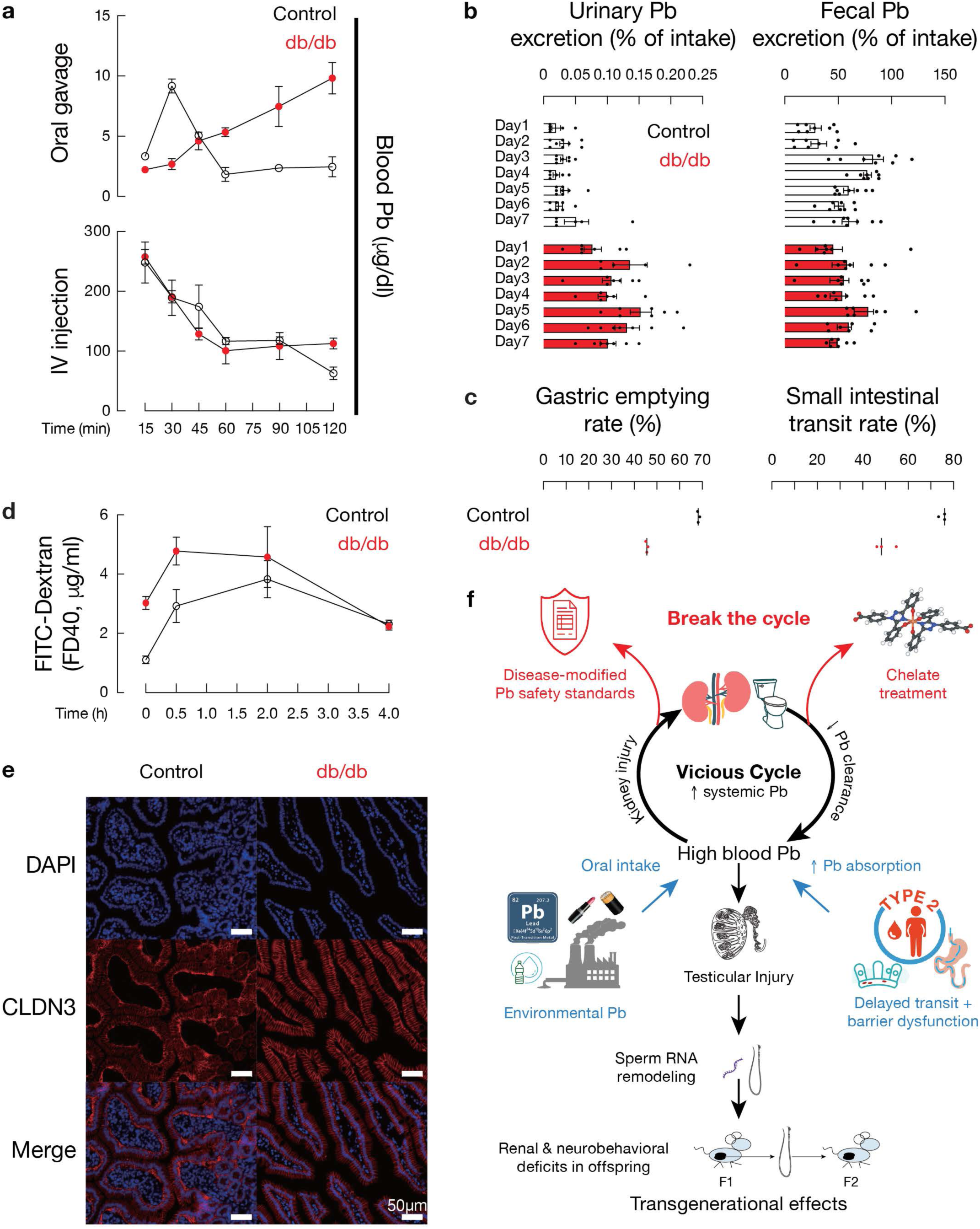
Diabetes slows gastrointestinal transit and enhances oral Pb absorption. **a**, Blood Pb concentration over time following oral gavage or intravenous injection of 2 mg kg⁻¹ Pb acetate in m/m and db/db mice (n = 3 per group). **b**, Proportion of total ingested Pb excreted in urine (left) and feces (right) relative to total Pb intake during the same period. **c**, Gastric emptying rate (phenol red recovery at 15 min) and small intestinal transit index. **d**, Plasma concentration of 40 kDa FITC–dextran (FD40) at 5 min, 30 min, 2 h and 4 h after gavage (44 mg per 100 g body weight). **e**, Representative confocal images of CLDN3 immunofluorescence (red) in duodenum of m/m and db/db mice. Nuclei stained with DAPI (blue). Scale bar, 50 μm. **f**, Conceptual model of diabetes-amplified Pb toxicity. Type 2 diabetes (T2D) increases intestinal Pb absorption through delayed gastrointestinal transit and epithelial barrier dysfunction, resulting in elevated systemic Pb burden. The kidney, a principal organ for Pb elimination, accumulates Pb and develops tubular injury, reducing clearance and further amplifying circulating Pb levels in a self-reinforcing cycle. Concurrently, increased Pb burden exacerbates testicular injury followed by transgenerational effects. These sperm RNA changes are sufficient to transmit renal and neurobehavioral phenotypes to F1 and F2 offspring. Potential intervention points include disease-informed Pb safety regulation and chelation therapy to reduce systemic Pb burden. Data are mean ± s.e.m.

Metabolic cage studies demonstrated that approximately half of ingested Pb was eliminated via feces in both genotypes (56 ± 26% in non-diabetic controls; 57 ± 24% in db/db mice, Fig. 3b, Extended Data Fig. 3b,c), consistent with established toxicokinetic evidence that fecal excretion represents the primary elimination route for ingested Pb, whereas urinary excretion accounts for a substantially smaller fraction^17,25^. When normalized to intake, the fraction of Pb eliminated in feces did not differ between genotypes. In contrast, absolute urinary Pb excretion was significantly higher in db/db mice (*P* = 1.69 × 10⁻¹⁶; Fig. 3b; Extended Data Fig. 3d), reflecting increased systemic Pb burden and compensatory renal elimination rather than impaired clearance capacity. The comparable proportional distribution of fecal and urinary excretion between groups indicates that diabetes does not alter the relative route of Pb elimination, but instead increases Pb absorption.

We next explored mechanisms underlying increased Pb absorption. Db/db mice exhibited delayed gastric emptying (*P* = 6.7 × 10⁻⁷) and reduced small-intestinal transit (*P* = 6.9 × 10⁻³) compared with non-diabetic controls (Fig. 3c), consistent with diabetic gastrointestinal dysmotility^26^. Prolonged luminal transit is expected to increase contact time between ingested Pb and the absorptive epithelium. To evaluate epithelial barrier properties, we assessed size-selective permeability using FITC–dextran tracers. Plasma fluorescence following 40 kDa FITC–dextran gavage was significantly elevated in db/db mice at early time points (5–120 min), whereas no difference was observed with 70 kDa FITC–dextran (Fig. 3d; Extended Data Fig. 3d). This pattern indicates increased permeability to intermediate-size molecules without evidence of gross barrier disruption. Immunofluorescence analysis of duodenal sections revealed altered junctional localization of CLDN3 (claudin-3) in db/db mice, with reduced apical enrichment compared with non-diabetic controls (Fig. 3e). CLDN3 is a tight junction component implicated in regulation of paracellular permeability^27,28^, and altered junctional organization has been reported in diabetic and obese models^29,30^. By contrast, ex vivo intestinal Pb uptake did not differ between genotypes (Extended Data Fig. 3e), arguing against enhanced transcellular Pb transport. Collectively, these findings indicate that delayed gastrointestinal transit and functional alterations in epithelial barrier properties act in concert to increase net oral Pb absorption in the diabetic state (Fig. 3f). Both mechanisms have parallels in human T2D^31,32^.

### Pb accelerates diabetic kidney injury and chelation improves renal outcomes

Given the strong association between elevated blood Pb and impaired renal function in individuals with T2D (Fig. 1b), we next examined whether diabetes-enhanced Pb burden exacerbates kidney injury in vivo. After 8 weeks of exposure, tissue distribution analysis revealed that the kidney was the second-largest Pb reservoir after bone (Extended Data Fig. 4a), consistent with its dual role as the route of Pb elimination and a major site of Pb accumulation^33,34^. Renal Pb concentrations reached 4.6 ± 2.4 μg g⁻¹ in db/db mice and 2.0 ± 0.2 μg g⁻¹ in non-diabetic controls, corresponding to 14- and 18-fold higher levels than blood, respectively. Longitudinal metabolic-cage analysis demonstrated that the blood-to-urine Pb ratio remained stable in non-diabetic controls but increased progressively in db/db mice over time (linear mixed-effects model, *P* = 1.4 × 10⁻³; Fig. 4a), indicating declining efficiency of renal Pb elimination in the diabetic state. The combination of increased renal Pb accumulation and reduced handling efficiency is consistant with progressive impairment of Pb clearance under sustained systemic burden.

**Figure 4.**
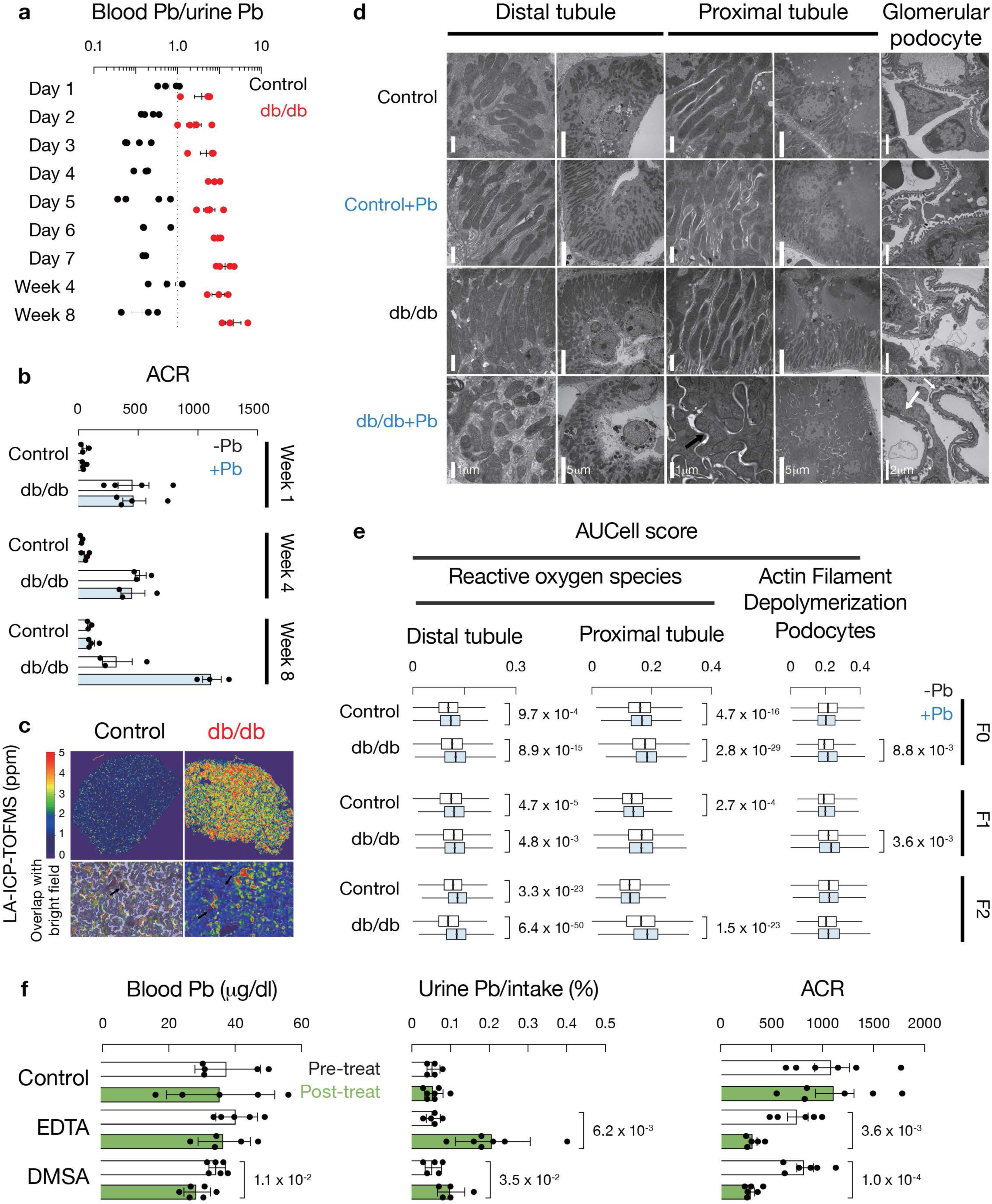
Chronic Pb exacerbates diabetic kidney injury and chelation improves renal outcomes. **a**, Ratio of blood Pb to urinary Pb concentration in m/m(Pb) and db/db(Pb) mice during days 1–7 and at weeks 4 and 8 (n = 3-4 per group). **b**, Urinary albumin-to-creatinine ratio (ACR) at weeks 1, 4 and 8. **c**, Laser ablation–ICP–MS heat map of Pb distribution in kidney sections. Representative images from 3 mice per group. **d**, Transmission electron micrographs of glomerular podocytes and tubular mitochondria after 8 weeks of Pb exposure. Representative images from 3 mice per group. Scale bars as indicated. **e**, AUCell pathway activity scores for reactive oxygen species (ROS) in proximal tubule (PT) and distal convoluted tubule (DCT) cells, and actin filament depolymerization in podocytes. **f**, Chelation therapy in db/db(Pb) mice. Blood Pb, urinary Pb excretion fraction, and ACR measured before and after 5-day treatment with EDTA (i.p.) or DMSA (oral). Data are mean ± s.e.m. Statistical tests as indicated.

To assess renal injury, we quantified urine microalbumin and calculated the albumin-to-creatinine ratio (ACR). Urine microalbumin reflects albumin excretion, whereas ACR normalizes for urine concentration variability and is the clinically preferred metric for detecting early glomerular injury^35^. Both urine microalbumin and ACR were significantly elevated in db/db mice after 8 weeks of Pb exposure (*P* = 2.1 × 10⁻² and *P* = 5.7 × 10⁻³, respectively), but not at weeks 1 or 4 (Fig. 4b; Extended Data Fig. 4b). The time-dependent emergence of albuminuria paralleled the progressive rise in blood-to-urine Pb ratio (Fig. 4a), consistent with a cumulative effect of retained Pb on renal function, and a progressive impairment of Pb clearance.

The observed ∼50–100% increase in ACR falls within the range reported in early-stage db/db progression models^36^. Structural injury remained limited: Hematoxylin and eosin staining revealed no overt morphological abnormalities (Extended Data Fig. 4c). Masson’s trichrome staining showed mild fibrosis in Pb-exposed non-diabetic controls (*P* = 0.030), whereas db/db mice exhibited elevated baseline fibrosis consistent with diabetic kidney disease. Importantly, db/db(Pb) mice did not display further fibrotic expansion relative to db/db controls (*P* = 0.80) (Extended Data Fig. 4d,e). Serum creatinine, blood urea nitrogen and estimated filtration indices remained within normal ranges (Extended Data Fig. 4f), a pattern characteristic of early-stage diabetic kidney injury in this model^36^.

Laser ablation–ICP–MS revealed preferential Pb enrichment in tubular regions outside glomeruli (Fig. 4c). Transmission electron microscopy demonstrated podocyte foot-process effacement and marked mitochondrial swelling in tubular epithelial cells of db/db(Pb) kidneys (Fig. 4d), consistent with proximal tubular Pb toxicity^15,33^. Single-cell RNA sequencing further showed that Pb exposure selectively upregulated podocyte remodeling programs in db/db mice and enriched reactive oxygen species (ROS) pathways in distal convoluted tubule (DCT) cells (Fig. 4e, Extended Data Fig. 4g.h). These findings implicate oxidative stress–mediated tubular injury as a key driver of diabetic Pb toxicity.

Importantly, the synergistic toxicity of diabetes and Pb was organ-specific. Liver enzymes, lipid profiles, and histology were unchanged by Pb exposure in both genotypes (Extended Data Fig. 4i–j). Echocardiography, retinal imaging, hematologic parameters, and systemic stress markers likewise showed no overt Pb-induced pathology beyond modest diabetic changes (Extended Data Fig. 4l–r).

Given prior clinical reports of slowed renal decline following EDTA chelation in selected patients with T2D and elevated Pb burden^10,11^, we tested whether reducing Pb burden mitigates renal injury in vivo. After 8 weeks of Pb exposure, db/db(Pb) mice were treated with EDTA or DMSA for 5 days. DMSA (succimer), an orally bioavailable chelator approved for pediatric Pb poisoning^37,38^, significantly reduced circulating Pb concentrations, whereas EDTA did not produce a significant short-term decline (Fig. 4f). Both chelators increased urinary Pb excretion and were associated with reductions in albuminuria and ACR (Fig. 4f), without overt toxicity (Extended Data Fig. 4s–u). Although EDTA did not measurably reduce circulating Pb during this short interval, increased urinary Pb excretion is consistent with mobilization of bioavailable Pb pools. These findings indicate that modifying Pb burden can partially attenuate renal stress even after chronic exposure.

### Pb exacerbates diabetes-associated testicular injury

Population-level analyses indicate that approximately 9–10% of men with T2D are of reproductive age^39^. In our cohort, men aged 18–45 years with T2D exhibited significantly higher blood Pb concentrations compared with age-matched non-diabetic controls (Fig. 5a). Pb has long been associated with impaired male fertility, including reduced sperm count, decreased motility and endocrine disruption^40–42^. We therefore asked whether diabetes modifies reproductive susceptibility to Pb exposure.

**Figure 5.**
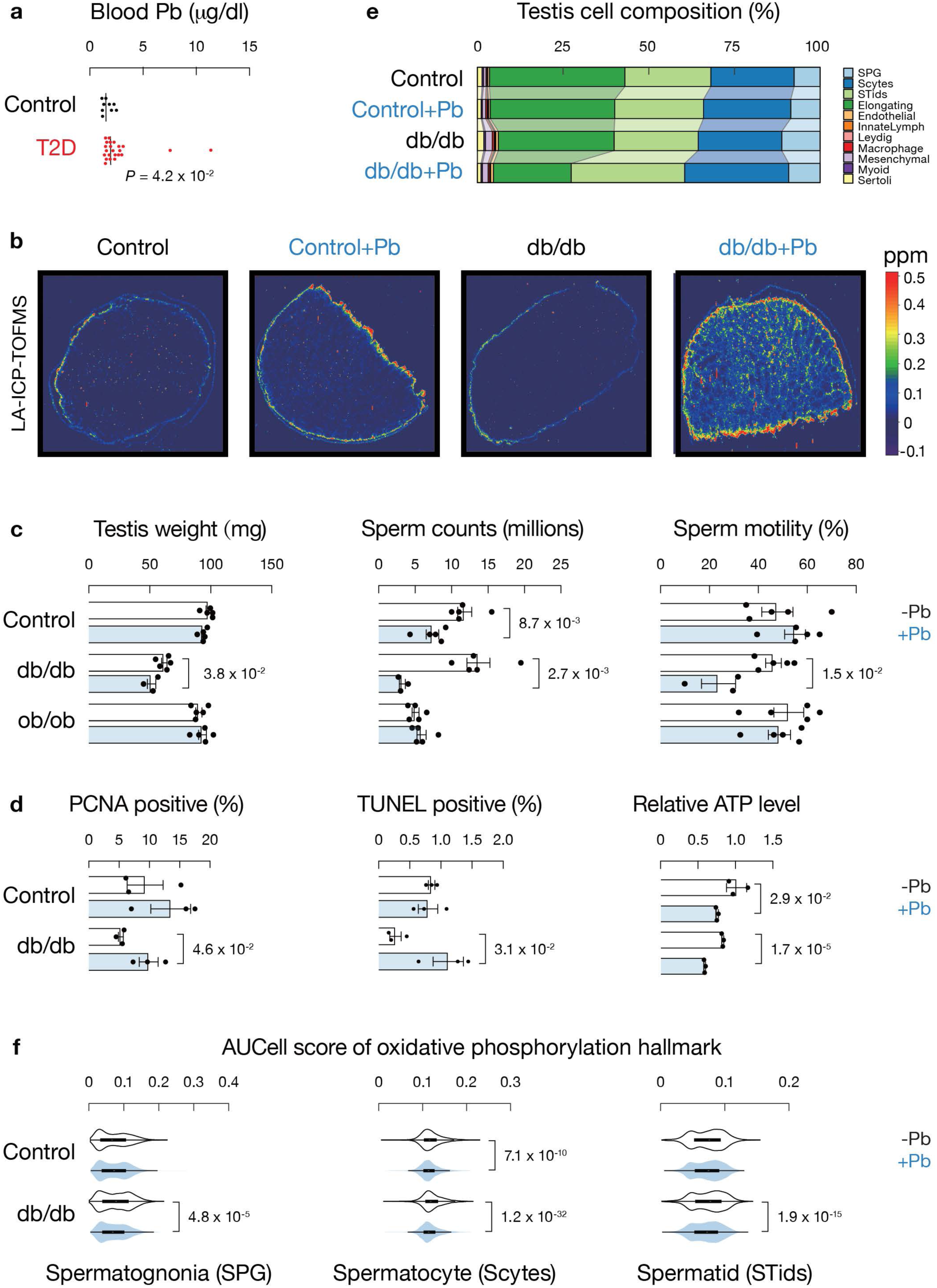
Pb exacerbates testicular dysfunction in diabetes. **a**, Blood Pb concentrations in men aged 18–45 years with and without T2D. **b**, Testicular Pb content measured by ICP–MS after 8 weeks of exposure. **c**, Testis weight, epididymal sperm count, and percentage of motile sperm in m/m, m/m(Pb), db/db, and db/db(Pb) mice (n = 3-5 per group). **d**, Quantification of PCNA-positive spermatogonia and TUNEL-positive germ cells; testicular ATP content normalized to total protein. **e**, Relative proportions of spermatogenic cell populations from scRNA-seq. **f**, AUCell activity scores for oxidative phosphorylation in spermatogonia (SPG), spermatocytes (Scytes), and round spermatids (STids). Data are mean ± s.e.m. Zhang et al, Fig. 6

Although the testis is not a major Pb storage organ compared with bone or kidney (Extended Data Fig. 4a), testicular Pb concentrations were increased 6.7-fold in db/db mice relative to non-diabetic controls (control: 8.9 ± 1.7 ng g⁻¹; db/db: 60 ± 41 ng g⁻¹; P = 0.049). Laser ablation–ICP–MS mapping demonstrated diffuse Pb distribution throughout the testicular parenchyma (Fig. 5b), consistent with penetration of the blood–testis barrier. Chronic Pb exposure significantly reduced testis weight, seminiferous tubule diameter, sperm count and sperm motility in db/db mice (Fig. 5c; Extended Data Fig. 5a). In non-diabetic controls, a modest reduction in sperm count was observed, but the magnitude of impairment was substantially greater in db/db mice. Obese but non-diabetic ob/ob mice did not exhibit similar exacerbation (Fig. 5c), indicating that the interaction between Pb and reproductive injury is specific to the diabetic metabolic state rather than obesity alone.

To define the cellular basis of reduced sperm output, we compared db/db mice with and without Pb exposure. Immunostaining for proliferating cell nuclear antigen (PCNA) revealed no reduction in spermatogonial proliferation in db/db mice following Pb exposure (Fig. 5d, left; Extended Data Fig. 5b). In contrast, TUNEL staining demonstrated a marked increase in germ-cell apoptosis in db/db(Pb) testes relative to db/db controls (Fig. 5d, middle; Extended Data Fig. 5c,d), consistent with seminiferous tubule atrophy. Single-cell RNA sequencing of whole testes further resolved stage-specific defects. A total of 277,421 cells were retained and classified into spermatogonia, spermatocytes, round spermatids and elongating spermatids, along with major somatic cell types (Extended Data Fig. 5e). In db/db mice exposed to Pb, the proportion of elongating spermatids decreased by 11.1%, whereas round spermatids increased by 8.6% compared with db/db controls (χ² test *P* = 1.2 × 10⁻^177^), indicating impaired transition from round to elongating spermatids (Fig. 5e). Consistent with PCNA staining, cell-cycle and DNA replication programs remained preserved in spermatogonia (Extended Data Fig. 5f). By contrast, apoptosis- and DNA damage–response pathways were preferentially upregulated in spermatocytes and spermatids in db/db(Pb) testes (Extended Data Fig. 5g). At the metabolic level, oxidative phosphorylation and mitochondrial energy-production pathways were significantly downregulated across spermatogonia, spermatocytes and round spermatids in db/db(Pb) mice relative to db/db controls (Fig. 5f). These transcriptomic alterations corresponded with reduced testicular ATP content (Fig. 5d, right) and are consistent with established mitochondrial toxicity of Pb^43^. Collectively, these data indicate that in the diabetic background, Pb exposure primarily enhances germ-cell apoptosis and disrupts spermatid maturation, rather than suppressing spermatogonial proliferation.

### Paternal diabetes and Pb exposure program heritable renal and neurobehavioral phenotypes via sperm RNA

To determine whether combined paternal diabetes and Pb exposure generate heritable phenotypes, we produced offspring by in vitro fertilization (IVF) using sperm from four paternal groups (non-diabetic controls, control(Pb), db/db and db/db(Pb)) to fertilize uniform oocytes from non-diabetic db/m females, followed by transfer into surrogate dams (Fig. 6a). IVF was used to eliminate potential confounding effects of seminal plasma, mating behavior and maternal metabolic status. F₁ males were subsequently used to generate F₂ offspring by the same IVF strategy, ensuring that F₂ animals were not directly exposed to paternal Pb. Persistence of phenotypes into F₂ in paternal exposure paradigms is considered consistent with transgenerational inheritance^44–46^.

**Figure 6.**
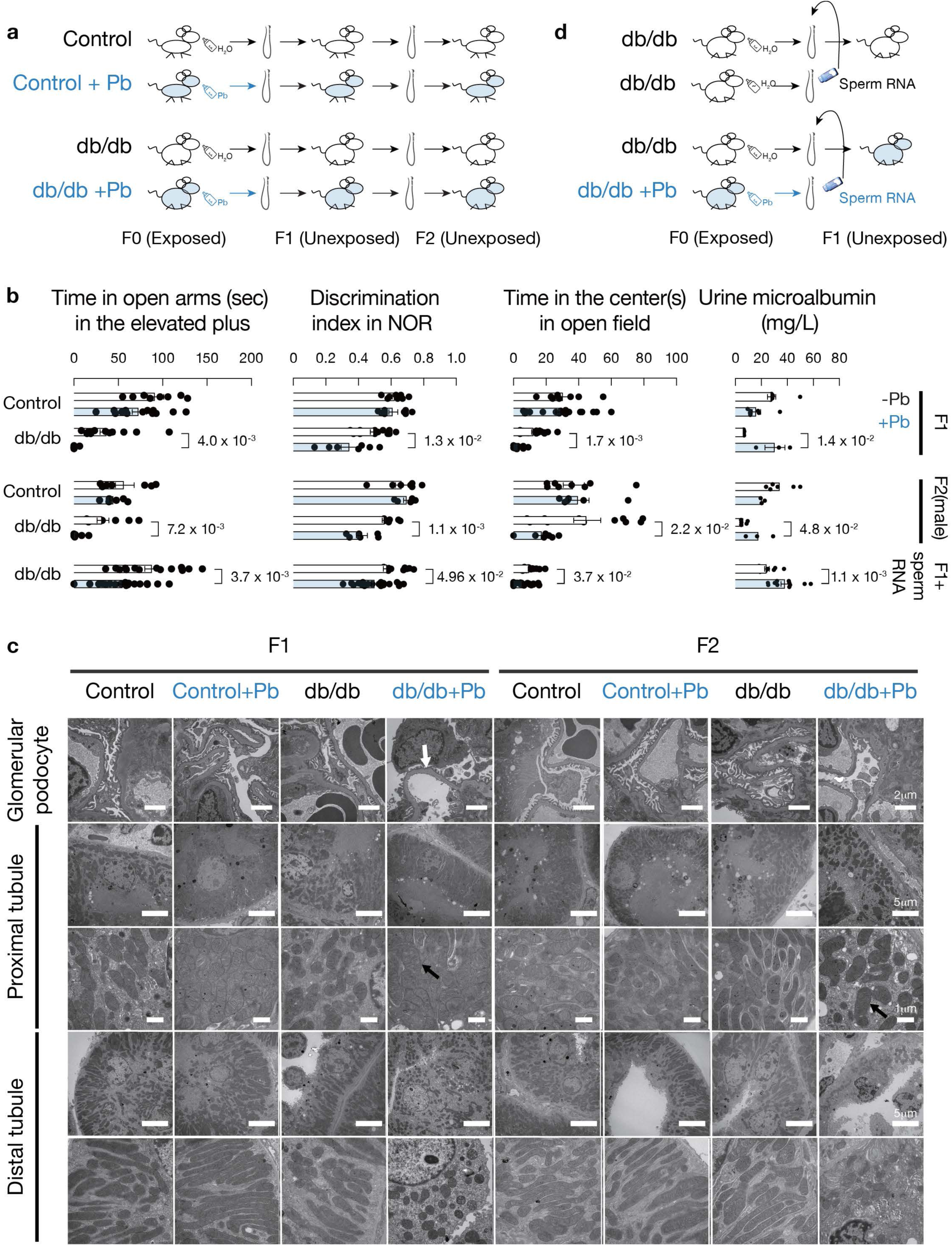
Paternal diabetes and Pb exposure induce transgenerational renal and neurobehavioral phenotypes. **a**, IVF breeding scheme for F₁ and F₂ generations. Sperm from m/m, m/m(Pb), db/db and db/db(Pb) males were used to fertilize wild-type oocytes. **b**, Behavioral assays (elevated plus maze, novel object recognition, open field) and urine microalbumin in F₁(total), F₂ (male)and F₁-injection(total) offspring. **c**, Transmission electron micrographs of kidneys from F₁ and F₂ offspring. Representative images of podocyte foot processes and tubular mitochondria. **d,** Microinjection scheme for F1-injection offspring. Sperm RNA was isolated from lead-exposed and unexposed db/db mice, and then was subsequently injected into the male pronucleus of normally fertilized zygotes to produce the F1-injection generation. Data are mean ± s.e.m. Statistical tests as indicated.

Behavioral phenotyping revealed metabolic context– and sex-dependent effects of paternal Pb exposure across domains of learning and emotional regulation. Spatial and recognition memory were assessed using the novel position recognition (NPR) and novel object recognition (NOR) paradigms. In the NPR task, paternal Pb exposure in non-diabetic control males was sufficient to induce impaired spatial recognition in F₁ offspring (total and male), and this deficit persisted into F₂ (total and male) (Extended Data Fig. 6a). In contrast, NPR impairment was not observed in the db/db background, indicating that metabolic context modifies the neural circuits susceptible to paternal Pb exposure. In the NOR assay, memory deficits were most pronounced in female F₁ and male F₂ offspring, demonstrating domain- and sex-specific penetrance.

Emotional behavior was assessed using the elevated plus maze and open-field assays, which measure anxiety-like exploration under conflict conditions. Offspring of db/db(Pb) sires exhibited increased anxiety-like behavior in the elevated plus maze in both male and female F₁ animals, with persistence into F₂, particularly in males (Fig. 6b; Extended Data Fig. 6b). In the open-field test, alterations were detected in F₁ offspring overall and in male F₁ and F₂ subsets. Locomotor capacity, assessed by treadmill performance, remained preserved (Extended Data Fig. 6c), excluding nonspecific motor impairment. Because F₂ female numbers were limited, F₂ analyses are presented primarily for males, with consistent directional trends observed in combined cohorts. Together, these findings demonstrate that adult paternal Pb exposure is sufficient to induce stable, heritable neurobehavioral alterations, and that diabetic metabolic state reshapes both the penetrance and neural circuitry underlying these effects. The observed sex-biased patterns are consistent with prior reports of sexually dimorphic inheritance following paternal environmental exposures^18,19^.

Unexpectedly, paternal diabetes and Pb exposure also programmed renal susceptibility across generations. Although gross renal morphology and fibrosis were unchanged in F₁ and F₂ offspring (Extended Data Fig. 6d, e), urine microalbumin and ACR were elevated in F₁, and urine microalbumin remained elevated in F₂ (Fig. 6b, right; Extended Data Fig. 6f), indicating persistent renal stress. Ultrastructural analysis revealed podocyte abnormalities and mitochondrial alterations in tubular epithelial cells in both generations (Fig. 6c). Single-cell RNA sequencing further demonstrated sustained activation of podocyte remodeling–associated programs in F₁ offspring and enrichment of reactive oxygen species (ROS) pathways in distal convoluted tubule (DCT) cells in both F₁ and F₂ descendants (Fig. 4e), consistent with inherited renal vulnerability.

Importantly, the heritable phenotypes were organ-specific rather than reflecting generalized developmental toxicity. No abnormalities were detected in testis weight, sperm count, sperm motility, liver histology or circulating metabolic markers in F₁ and F2 offspring (Extended Data Fig. 6g–i), indicating that paternal exposure did not induce systemic developmental impairment. The behavioral abnormalities occurred in the absence of motor deficits, and the renal phenotype was characterized by stress markers and ultrastructural changes without overt fibrosis, consistent with increased susceptibility rather than terminal pathology.

To test whether sperm-borne RNA is sufficient to transmit these phenotypes, we microinjected total sperm RNA isolated from db/db(Pb) males or db/db controls into naïve zygotes and phenotyped the resulting offspring (Fig. 6d). Offspring derived from db/db(Pb) sperm RNA recapitulated anxiety-like behavior, recognition-memory deficits and renal stress, including elevated urine microalbumin and ACR (Fig. 6b; Extended Data Fig. 6b,f). This experimental paradigm parallels established demonstrations that sperm RNA is sufficient to transmit paternal exposure phenotypes when introduced into zygotes^18–21^. Collectively, these findings indicate that combined paternal diabetes and Pb exposure program heritable renal and neurobehavioral phenotypes and that sperm RNA is sufficient to mediate this transmission.

### Trace Pb is detectable in consumer products relevant to oral exposure

The enhanced oral Pb absorption observed in diabetic mice prompted us to assess whether trace Pb in commonly consumed products could plausibly contribute to systemic burden. To provide environmental context for the human cohort, we measured Pb concentrations in drinking water from three municipal reservoirs supplying the study region; Pb levels were below the detection limit in all samples (Extended Data Table 1). We then quantified Pb in three independent batches of ten widely consumed sugar-free beverages and in lip products from ten leading brands. Detectable but low Pb concentrations were observed in beverages (maximum 0.77 ± 0.07 μg L⁻¹) and lip products (maximum 0.14 μg g⁻¹) (Extended Data Table 1). All measured values were substantially below current regulatory action levels in both the United States and China.

To estimate exposure magnitude, we combined measured concentrations with published consumption data. At the highest detected level (0.77 μg L⁻¹), consumption of 2 L per day would correspond to approximately 1.5 μg Pb daily from beverages. In contrast, average lip product use (24 mg per day) would yield ∼0.003 μg Pb at the maximum measured concentration. Thus, among the products examined, beverages represent a quantitatively greater oral exposure source under typical use. Although well below regulatory action levels, chronic low-level intake may incrementally increase systemic Pb burden. In T2D, where intestinal absorption is enhanced and renal clearance efficiency declines, even modest sustained exposure could have disproportionate biological consequences.

## Discussion

This study reframes the long-debated relationship between Pb exposure and diabetes by demonstrating that metabolic disease modifies the physiological interfaces governing toxicant handling. In diabetic mice, delayed gastrointestinal transit and altered epithelial barrier properties increased oral Pb absorption without impairing systemic elimination at early stages. With sustained exposure, however, the kidney—a principal organ for Pb accumulation and excretion—developed enhanced injury under elevated Pb burden. Even modest tubular dysfunction reduced effective Pb clearance, establishing a self-reinforcing cycle in which diabetes promotes Pb retention and retained Pb further stresses renal tissue (Fig. 3f).

Although T2D is strongly linked to diabetic kidney disease (DKD), it does not fully explain the marked heterogeneity in DKD onset and progression^47,48^. Classical rodent models, including db/db mice, typically exhibit early mesangial expansion and mild albuminuria but rarely progress to advanced glomerulosclerosis or renal failure without additional genetic or environmental stressors^49^. In contrast to the controlled conditions of laboratory housing, human populations are continuously exposed to low-level environmental toxicants. By introducing chronic, low-level Pb exposure into db/db mice, we uncovered a nephrotoxic interaction that accelerated renal injury specifically within the diabetic context. These findings position Pb not merely as a background exposure, but as a context-dependent modifier of DKD trajectory, offering a mechanistic explanation for part of the unexplained variability in diabetic renal outcomes.

Our identification of Pb as a contributor to a self-reinforcing cycle of renal stress, together with the observation that short-course chelation attenuated renal stress markers following chronic Pb exposure in the mouse model, suggests that reducing Pb burden may slow disease progression. This interpretation aligns with randomized clinical studies in which calcium disodium EDTA reduced the rate of GFR decline in patients with diabetic nephropathy and elevated body Pb burden^10,11^. Chelation therapy is an established intervention for clinically elevated Pb levels; what remains uncommon is systematic assessment of Pb burden in patients with DKD. Standard DKD management emphasizes glycaemic control and renoprotective pharmacotherapy, yet substantial residual risk persists despite RAAS blockade, SGLT2 inhibitors, GLP-1 receptor agonists and mineralocorticoid receptor antagonists^50,51^. Together, our findings and prior clinical evidence suggest that identifying and interrupting Pb-driven amplification of renal stress may represent a tractable and currently under-recognized therapeutic axis in DKD, warranting prospective evaluation in carefully stratified patient populations.

Beyond somatic organ injury, our findings provide experimental evidence that a common metabolic disorder can synergize with an environmental toxicant to induce heritable renal susceptibility. Unlike prior reports in which transgenerational kidney phenotypes were associated primarily with endocrine-disrupting chemicals^52,53^, neither T2D alone^54^ nor Pb exposure alone under comparable conditions^5^ produced persistent renal dysfunction across generations. These results support a disease–toxicant interaction model in which metabolic state reshapes germline information and modulates offspring organ vulnerability. Given the global prevalence of both T2D and environmental Pb exposure^55^, this work introduces a reproductive dimension to metabolic disease, suggesting that parental health and exposures before conception may influence renal risk in descendants through sperm-mediated epigenetic mechanisms.

At the level of environmental health policy, our results highlight a structural limitation of existing Pb safety standards, which are largely derived from general-population risk models and implicitly assume uniform susceptibility. By demonstrating that T2D alters Pb toxicokinetics and amplifies toxicity at exposure levels currently deemed acceptable, our findings argue for a disease-informed approach to risk assessment. Just as children are recognized as uniquely vulnerable to Pb neurotoxicity, chronic metabolic diseases such as T2D may constitute adult vulnerability states that warrant distinct consideration. Incorporating disease status into exposure guidelines—whether through targeted monitoring, refined safety margins or stratified regulatory thresholds—could enhance protection for populations disproportionately affected by metabolic disease. More broadly, recognizing disease-modified toxicokinetics as a regulatory principle may be essential for equitable public health protection in an era of widespread diabetes.

In summary, we identify a previously unrecognized vicious cycle linking T2D and environmental Pb exposure and redefine Pb as a context-dependent amplifier of disease. These findings establish disease-modified toxicokinetics as a fundamental mechanism linking metabolic disease, environmental exposure and inherited vulnerability, with implications for DKD management and environmental health policy.

## Methods

### Human participants and study design

This retrospective age- and sex-matched case–control study included 118 individuals with type 2 diabetes mellitus (T2D) and 61 non-diabetic controls recruited in 2023 at the Fourth Affiliated Hospital, Zhejiang University School of Medicine (Yiwu, China). Participants were aged 26–79 years (mean ± s.d. = 52.07 ± 13.30 years). Diabetes subtype assignment (T2D, type 1 diabetes, juvenile-onset diabetes, gestational diabetes) was based on medical records and standard clinical diagnostic criteria used at the hospital^56^. Controls were recruited from the hospital Physical Examination Center and had no known history of diabetes, with fasting glucose < 5.6 mmol L⁻¹ and HbA1c < 5.7% (institutional reference ranges).

Exclusion criteria included known occupational Pb exposure, acute illness at the time of sampling, and chronic conditions expected to substantially alter Pb kinetics or metabolic indices (including malignancy, chronic liver disease, non-diabetic chronic kidney disease, and endocrine/metabolic disorders affecting glucose homeostasis). Participants using chelating agents at the time of sampling were excluded. The study protocol was approved by the Clinical Research Ethics Board of the Fourth Affiliated Hospital of Zhejiang University School of Medicine (Approval ID: K2023170) and conducted in accordance with the Declaration of Helsinki. Written informed consent was waived under ethics approval for retrospective analysis.

### Clinical chemistry and eGFR calculation

Serum indices—including alanine aminotransferase (ALT), aspartate aminotransferase (AST), alkaline phosphatase, total bilirubin, blood urea nitrogen (BUN), creatinine, electrolytes, fasting glucose, triglycerides, total cholesterol, and LDL—were measured using an automated chemistry analyzer (Hitachi Labospect 008 AS) according to the manufacturer’s instructions. Urine indices (including microalbumin and creatinine) were measured using an automated analyzer (Hitachi 3500) according to manufacturer protocols. Laboratory staff were blinded to case/control status. Estimated glomerular filtration rate (eGFR) was calculated using the CKD-EPI equation (2021 version) with creatinine in μmol L⁻¹ and unit conversion as specified by the equation implementation^57^. The urinary albumin-to-creatinine ratio (ACR) was calculated as: ACR (mg g⁻¹) = [urinary albumin (mg L⁻¹) ÷ urinary creatinine (μmol L⁻¹)] × 8,840.

### Statistical analysis for the human cohort

Normality was assessed using the Shapiro–Wilk test. Because blood Pb levels were non-normally distributed in the T2D cohort, group comparisons were performed using Mann–Whitney U tests unless stated otherwise. Correlations between blood Pb and clinical indices were assessed using Spearman’s rank correlation (two-tailed). Multiple testing correction for multi-element analyses and correlation panels used Benjamini–Hochberg false discovery rate control (FDR threshold 0.05), where applicable.

### Pb and multi-element quantification by ICP–MS

Whole-blood and tissue Pb concentrations were quantified by inductively coupled plasma–mass spectrometry (ICP–MS) using an Agilent 7800 quadrupole ICP–MS equipped with a collision/reaction cell operated in helium collision mode to reduce polyatomic interferences.

For whole blood, 100 μL of sample was digested in acid-cleaned PTFE tubes following a published workflow^58^. Briefly, 0.50 mL of concentrated nitric acid (65% HNO₃; Merck, Cat# 1.00456.2508) was added to each sample and incubated at 100 °C for 60 min. After cooling, digests were diluted to 3.0 mL with ultrapure water (18.2 MΩ·cm) and filtered through a 0.22 μm hydrophilic PTFE syringe filter (Anpel, Cat# SCAA-114). For solid tissues (kidney, bone, testis, liver, and other organs as indicated), approximately 0.1 g (wet weight) was digested in 1 mL nitric acid using the same conditions and brought to a final volume of 5 mL with ultrapure water. Procedural blanks and matrix-spiked recoveries were processed alongside samples.

ICP–MS was tuned daily. Typical operating parameters were RF power 1550 W, plasma gas flow 15 L min⁻¹, auxiliary gas 1.0 L min⁻¹, nebulizer gas 1.05 L min⁻¹, and helium collision gas flow ∼5.0 mL min⁻¹. Samples were introduced via autosampler with an integration time of 0.3 s and three technical replicate measurements per sample. Calibration was performed using NIST-traceable multi-element standards across three calibration points, prepared in matrix-matched acid diluent. Bismuth(1 μg ml^-1^) was used as an internal standard. Limits of detection (LOD) and quantification (LOQ) were defined as 3× and 10× the standard deviation of procedural blanks, respectively (blood Pb LOD = 0.03 μg L⁻¹; LOQ = 0.1 μg L⁻¹). Accuracy was verified using certified reference material NIST SRM 955c (Toxic Metals in Caprine Blood).

For exploratory multi-element profiling, a subset of human samples (controls n = 6; T2D n = 24) was analyzed using the same ICP–MS platform and digestion workflow. Of 27 candidate elements screened, only 15 elements above LOQ in ≥80% of samples were included in downstream statistical comparisons (Extended Data Table 1).

### Environmental and consumer product Pb measurements

To assess potential low-level exposure sources, three independent batches of 10 sugar-free beverages and 10 lipstick products (brands anonymized) were purchased in Yiwu, China and analyzed by ICP–MS. For beverages, 500 μL of product was mixed with 1.5 mL dilute nitric acid (2%) prior to analysis. For lipsticks, 0.1 g of homogenized product was digested in 3 mL nitric acid at 100 °C for 2 h and diluted to 5 mL before ICP–MS measurement. To exclude localized water contamination as a confounder for the Yiwu cohort, drinking water was collected from three municipal reservoirs (Suxi, Nanshan, Xianding) using trace-metal clean procedures (acid-washed containers; field blanks: ddH₂O) and analyzed by ICP–MS. Pb was below detection limit in all samples (LOD = 0.096 μg L⁻¹; reported in Extended Data Table 1).

### Mouse studies

#### Mouse models, husbandry, and experimental design

All animal procedures were approved by the Zhejiang University Laboratory Animal Welfare and Ethics Committee (Approval No. ZJU20250897) and performed in compliance with institutional guidelines. Mice were housed under specific pathogen-free conditions at 20–26 °C and 40–70% relative humidity on a 12 h light/12 h dark cycle (lights on 07:30–19:30), with ad libitum access to standard chow and water except during designated fasting periods. Cage density was five mice per cage (db/db mice: three per cage). Animals were randomized to experimental groups using computer-generated randomization, and outcome assessors were blinded to group allocation where feasible (allocation codes maintained by an independent individual; analyses performed on coded data). Mouse strains included leptin receptor mutant db/db mice on a C57BLKS/J background (GemPharmatech, stock no. T002407), leptin knockout ob/ob mice on a C57BL/6J background (GemPharmatech, stock no. T001461), and age-matched **non-diabetic controls (m/m)** (referred to as “non-diabetic controls” throughout the manuscript). Unless otherwise specified, adult male mice were used and experiments began at ∼8 weeks of age.

#### Pb exposure and chelation regimens

For chronic exposure, mice received drinking water containing 100ppm lead (Pb) acetate (approximately 63 ppm Pb) or control water for 8 weeks. Lead(II) acetate was selected because of its high aqueous solubility and established use in experimental Pb toxicology models^5,13,14,25^. Although environmental Pb occurs in multiple chemical forms (e.g., Pb sulfide, Pb oxide, Pb chromate) with differing solubility and bioavailability, the acetate form provides a reproducible and bioavailable exposure paradigm for mechanistic studies. Whether diabetes differentially alters absorption of less soluble Pb species remains to be determined.

Water intake was recorded at 24 h intervals. Blood Pb was measured at 14 time points (Day 1–6 and weeks 1–8). For controlled intake experiments, mice received a single oral gavage of lead(II) acetate (dose 2 mg Pb kg⁻¹), and Pb excretion in urine and feces was quantified over 24 h. For chelation experiments, db/db(Pb) mice were treated after 8 weeks of exposure with calcium disodium EDTA (150 mg kg⁻¹; intraperitoneal; once daily for 5 days) or succimer (DMSA; 50 mg kg⁻¹; oral; once daily for 5 days). Body weight and renal injury readouts (urinary albumin and creatinine for ACR calculation) were monitored during treatment.

#### Glucose and insulin tolerance testing

Intraperitoneal glucose tolerance tests (GTT) and insulin tolerance tests (ITT) were performed at week 8 of exposure following established protocols^59,60^. For GTT, mice were fasted for 16 h and injected intraperitoneally with D-glucose (2 g kg⁻¹; Sigma, Cat# G8769). For ITT, mice were fasted for 4 h and injected intraperitoneally with insulin (1.0 U kg⁻¹; ProCell, Cat# PB180432). Tail-vein blood glucose was measured at 0, 15, 30, 60, 90, and 120 min using a glucometer (Accu-Chek Performa, Roche). Area under the curve (AUC) was calculated using the trapezoidal method.

#### Pb pharmacokinetics

To compare oral versus systemic handling, mice received lead(II) acetate (2 mg Pb kg⁻¹) via oral gavage or tail-vein injection. Blood was collected at 15, 30, 45, 60, 90, and 120 min and Pb concentration measured by ICP–MS.

### Gastrointestinal assays

#### Gastric emptying and intestinal transit

Gastric emptying was quantified using the phenol red recovery method^61^. After a 4 h fast, mice were gavaged with 100 μL phenol red meal (0.5% phenol red in isotonic saline, pH 7.5). Fifteen minutes later, stomach contents were recovered and absorbance measured at 480 nm. Gastric emptying was calculated as 100 × [1 − (phenol red remaining/phenol red administered)]. Small intestinal transit was measured by charcoal meal test^61^. After a 4 h fast, mice were gavaged with 100 μL 10% activated charcoal and euthanized after 15 min. Transit index was calculated as (distance traveled/total small intestinal length) × 100%.

#### FITC–dextran permeability assay

Intestinal permeability was assessed using FITC–dextran tracers (40 kDa and 70 kDa; Sigma-Aldrich). After a 4 h fast, mice were gavaged with 44 mg per 100 g body weight (40 kDa) or 125 mg per 100 g body weight (70 kDa). Plasma was collected at 5 min, 30 min, 2 h, and 4 h (40 kDa), and at 30 min (70 kDa). Fluorescence was quantified using a plate reader (Biotek; excitation 480 nm, emission 520 nm) against standard curves^62^.

### Ex vivo uptake and tight junction imaging

#### Ex vivo intestinal Pb uptake

Intestinal segments (duodenum, jejunum, ileum, colon; 2 cm each) were isolated, rinsed, and incubated luminally with lead(II) acetate (100 ppm) for 30 min at 37 °C. Tissues were washed extensively, digested in nitric acid, and Pb content quantified by ICP–MS.

#### CLDN3 immunofluorescence

Duodenum tissues were fixed in 4% paraformaldehyde, cryoprotected, and cryosectioned at 7 μm. Sections were immunostained for claudin-3 (CLDN3; Thermo Fisher, Cat# 34-1700, 1:100) and imaged using identical acquisition settings across groups. Junctional localization was quantified in ImageJ/Fiji as mean junctional intensity per field (three fields per mouse).

#### Metabolic cage studies and renal function markers

Mice were acclimated to metabolic cages for 1 day and subsequently housed individually for 24 h for quantitative urine and feces collection. Water intake was recorded over the same period. Urine and fecal samples were digested in concentrated nitric acid, and Pb concentrations were quantified by ICP–MS as described above. Urine microalbumin and creatinine were measured using a Hitachi 3500 Fully Automated Biochemical Analyzer (Shanghai Bairong, Cat# E141332E; Wako Pure Chemical Industries, Cat# 509452) according to the manufacturers’ instructions. The albumin-to-creatinine ratio (ACR) was calculated to normalize for urine concentration variability. Serum creatinine and blood urea nitrogen (BUN) were measured using an automated chemistry analyzer (Hitachi Labospect 008 AS; FUJIFILM, Cat# 509452 and Cat# 507393) following manufacturer protocols.

#### Glomerular filtration rate (GFR) measurement

Transdermal GFR was measured using FITC–sinistrin clearance as previously described^63,64^. Briefly, mice were anesthetized with isoflurane for device placement. A miniaturized fluorescence detector (MediBeacon GmbH, Mannheim, Germany), consisting of light-emitting diodes, a photodiode, and an integrated battery, was affixed to the shaved and depilated dorsal skin using double-sided adhesive tape. Baseline background fluorescence was recorded for 5 min prior to intravenous injection of FITC–sinistrin (70 mg kg⁻¹; MediBeacon GmbH). After injection, fluorescence decay was recorded continuously for approximately 1.5–2.5 h while animals were awake and housed individually. The detector was then removed and data were analyzed using MB Lab/MB Studio software (MediBeacon GmbH). GFR (µL min⁻¹ or µL min⁻¹ per body weight) was calculated from the plasma elimination kinetics of FITC–sinistrin, derived from the exponential decay of fluorescence intensity over time. Clearance curves were fitted using a one-, two-, or three-compartment model as appropriate, incorporating body weight and an empirical conversion factor as described previously^64^.

#### Histology and immunostaining

Kidney and testis tissues were processed for hematoxylin & eosin (H&E) staining using standard paraffin embedding and sectioning (section thickness 3um). For kidney fibrosis, Masson’s trichrome staining was performed using a commercial kit (Beyotime, Cat# C0189S), and fibrotic area was quantified using ImageJ from three non-overlapping fields per section across three sections per animal.

For testis apoptosis, TUNEL staining was performed on cryosections using a commercial kit (Elabscience, Cat# E-CK-A321) following manufacturer instructions, with DNase-treated positive controls and enzyme-omitted negative controls^65^. For spermatogonial proliferation, PCNA immunostaining was performed using mouse anti-PCNA (Santa Cruz, Cat# SC-25280; 1:500) and appropriate secondary antibodies^66^. Positive cells were quantified in 3 seminiferous tubule cross-sections per animal using consistent thresholds.

#### Transmission electron microscopy

Kidney tissues were fixed in 2.5% glutaraldehyde in 0.1 M sodium cacodylate buffer (pH 7.4), post-fixed in 1% osmium tetroxide, dehydrated through graded ethanol, embedded in resin, sectioned at 110 nm, and stained with uranyl acetate and lead citrate. Samples were imaged on a Thermo Scientific Talos L120C TEM. Podocyte foot-process effacement was quantified as the percentage of capillary loop circumference with effaced foot processes and scored on a 0–3 scale. Mitochondria were classified as normal, swollen, elongated, or cristae-disrupted, and the fraction of each category was reported per field. Quantification was performed in 3 fields per sample at 5,300–8,500× magnification.

#### Laser ablation ICP–MS imaging

Fresh-frozen tissues were cryosectioned at 20 μm and analyzed by ICP–MS system (icpTOF R, TOFWERK) coupled to Laser Ablation system (Matrix-Array Femtosecond Laser Ablation SystemGenesisGEO, Shanghai Chemlab Instrument). Ablation parameters included wavelength 343 nm, spot size ∼15 × 15 μm^2^, ablation frequency 40 Hz, scan speed 600 μm s⁻¹, and no reaction gas. Pb distribution maps were reconstructed from 208Pb signal intensity.

### Single-cell RNA sequencing

#### Tissue dissociation and cell preparation

Single-cell RNA sequencing (scRNA-seq) was outsourced to two service providers. Testis samples were collected from F0 mice (non-diabetic controls, control(Pb), db/db, db/db(Pb)) and processed using the Majorbio workflow. Kidney samples were collected from F0, F1, and F2 mice. For F0 kidney, two biological replicates per group were analyzed: one replicate processed with Majorbio (four samples total) and one processed with Annoroad (four samples total). All F1 and F2 kidney samples (eight samples total) were processed using the Annoroad workflow.

Majorbio workflow: Tissues were rinsed in PBS, minced, and digested in 0.25% trypsin with DNase I (10 μg mL⁻¹) in PBS supplemented with 5% FBS at 37 °C with shaking (50 rpm) for ∼40 min. Dissociated cells were collected every 20 min. Suspensions were filtered (40 μm), red blood cells lysed, washed, and viability assessed by AO/PI staining. Annoroad workflow: After performing cardiac perfusion with pre-chilled PBS, the kidneys were harvested. Tissues were minced (∼1–2 mm³), digested in trypsin at 37 °C for 15–30 min with gentle trituration, neutralized with DMEM + FBS, filtered (70 μm then 40 μm), and centrifuged (500×g, 5 min, 4 °C). Quality-control criteria were viability >80%, aggregate rate <5%, debris <5%.

#### Library preparation and sequencing

Majorbio libraries were prepared using Chromium Single Cell 3′ v4.1 chemistry following manufacturer protocol and sequenced on Illumina NovaSeq X Plus with paired-end 150 bp reads. Annoroad libraries were prepared using Chromium GEM-X Single Cell 3′ Kit v4 (10x Genomics, PN-1000686) and Single Cell Chip (PN-1000690), targeting 20,000 recovered cells per sample and sequenced to ∼40,000 reads per cell on Illumina NovaSeq X-25B.

#### Computational analysis

Raw sequencing reads were aligned to the mouse reference genome (mm10) using Cell Ranger v9.0.1^67^. Ambient RNA contamination and background signal were removed using CellBender v0.3.0^68^ with a target false-positive rate of 0.01 and 150 training epochs. The learning rate was set to 6 × 10⁻⁶ for kidney datasets and 1 × 10⁻⁵ for testis datasets. Background-corrected count matrices were analyzed using Scanpy v1.11.4^69^.

Genes detected in fewer than three cells were excluded. Cells were removed if they met any of the following criteria: total unique molecular identifiers (UMIs) <500, fewer than 250 detected genes, mitochondrial transcript fraction >40%, or >80% contribution from the top 20 most highly expressed genes. For kidney datasets, cells with >5,000 detected genes were additionally excluded to reduce potential multiplets. Doublets were identified using scDblFinder with default parameters and removed prior to downstream analysis^70^.

Filtered count matrices were normalized to 10,000 counts per cell and log-transformed. Cell-cycle phase scores were calculated using curated S-phase and G2/M gene sets^71^. The top 3,000 highly variable genes were identified using the Seurat v3 method^72^. Batch effects were corrected using Harmony^73^. A k-nearest neighbor graph was constructed using 30 neighbors and the first 20 principal components. Two-dimensional visualization was performed using uniform manifold approximation and projection (UMAP) with a minimum distance of 0.2^74^. Clustering was conducted using the Leiden algorithm at a resolution of 0.1^75^.

Kidney cell types were annotated based on canonical marker genes, including proximal tubule cells (*Slc34a1*, *Lrp2*), podocytes (*Nphs1*, *Nphs2*, *Wt1*), loop of Henle (*Umod*, *Slc12a1*), distal convoluted tubule (*Slc12a3*), collecting duct principal cells (*Aqp2*), intercalated cells (*Atp6v1g3*, *Slc4a1*), endothelial cells (*Pecam1*, *Kdr*), fibroblast/mesenchymal-like cells (*Pdgfrb*, *Dcn*), myeloid cells (*Itgam*), T cells (*Cd3e*, *Cd3g*), natural killer (NK) cells (*Nkg7*, *Klrb1c*), B cells (*Cd19*), and plasma cells (*Sdc1*, *Jchain*). Testis cell types were annotated by reference mapping using scArches^76^ with scANVI^77^, using an adult mouse testis reference atlas^78^.

Gene set activity scores were computed using AUCell^79^. Group differences in AUCell scores were assessed using two-sided Wilcoxon rank-sum tests. Differential gene expression analysis was performed using MAST with batch included as a covariate^80^. Multiple testing correction was performed using the Benjamini–Hochberg false discovery rate method where applicable.

#### Reproductive phenotyping, testosterone, and ATP

Epididymal sperm were released from cauda epididymides into pre-warmed Whitten’s medium (37 °C) and counted using a hemocytometer. Motility was assessed by light microscopy at 37 °C and calculated as (motile sperm/total sperm) × 100%. Serum testosterone was measured using an ELISA kit (SIMIBIO, Cat# ME-M1435S) from morning samples (09:00–11:00). Testicular ATP was quantified using a luciferase-based ATP assay kit (Beyotime, Cat# S0027) and normalized to total protein measured by BCA assay (Beyotime, Cat# P0010).

#### IVF, embryo transfer, and multigenerational design

To generate F1 and F2 offspring, sperm from F0 males (non-diabetic controls, control(Pb), db/db, db/db(Pb)) was used for IVF with oocytes from unexposed wild-type females. Females were superovulated with PMSG (10 IU) followed 48 h later by hCG (10 IU). Cumulus–oocyte complexes were collected 14–16 h after hCG. Sperm were capacitated in HTF medium (Nanjing Aibei Biotechnology, Cat# M1135) and co-incubated with oocytes at 1 × 10⁶ mL⁻¹ sperm concentration for 5 h. Embryos were cultured to the 2-cell stage and transferred into day 0.5 pseudopregnant ICR surrogates (10–20 embryos per surrogate). F2 offspring were generated by repeating IVF using sperm from adult F1 males.

#### Sperm total RNA extraction and zygote microinjection

Total sperm RNA was isolated from cauda epididymal sperm of db/db(Pb) and db/db (without Pb) males as described previously^81^, with somatic lysis buffer to minimize somatic cell contamination, and DNase treatment (Turbo DNA-free kit, Cat# 3091668) to eliminate residual genomic DNA contamination. RNA concentration was quantified using a Qubit fluorometer (Thermo Fisher Scientific), and RNA integrity and size distribution were assessed using an Agilent 4150 TapeStation system.

As described previously^18^, purified sperm RNA was diluted to 2 ng μL⁻¹ in RNase-free injection buffer. Approximately 5–10 pL of total RNA was microinjected into the male pronucleus of fertilized zygotes at the one-cell stage using an Olympus IX73 inverted microscope equipped with an Eppendorf TransferMan 4R micromanipulator and FemtoJet 4i microinjection system. Successfully injected embryos were cultured in vitro to the 2-cell stage under standard conditions and subsequently transferred into the oviducts of day 0.5 pseudopregnant surrogate females for gestation.

### Behavioral testing

Behavioral assessments were conducted during the light phase (09:00–17:00) by experimenters blinded to genotype and treatment. Mice were acclimated to the testing room and experimenters for at least 7 days prior to behavioral experiments. The behavioral battery included open field, elevated plus maze, novel object recognition (NOR), and novel position recognition (NPR) tests^82–84^.

For the Novel Position Recognition (NPR) assay, mice were tested in a 40 × 40 cm open-field arena under uniform illumination. During the habituation phase, mice were allowed to explore two identical objects placed symmetrically in the arena for 10 minutes. After a 6-hour retention interval in the home cage, one object was displaced to a novel location while the other remained in its original position. Mice were reintroduced to the arena for a 10-minute test session. Behavior was video-recorded and analyzed using EthoVision XT (Noldus Information Technology). Object exploration was defined as direct nose-oriented investigation within close proximity (<2 cm). Quantification was performed over the last 5 minutes of the test session. The discrimination index was calculated as:

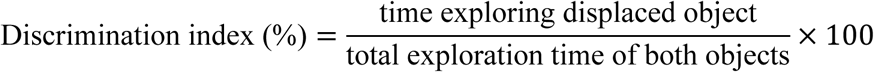

^85^.

Between trials, the apparatus and objects were cleaned with 70% ethanol to eliminate olfactory cues.

Treadmill endurance testing was performed using a graded running protocol following 5 days of acclimation (treadmill model SA101; manufacturer Science, China).

### Additional phenotyping

Echocardiography was performed at week 16 using Sigma Vet under 1–2% isoflurane anesthesia. LV dimensions and functional parameters were derived by M-mode analysis. SD-OCT and fundus imaging were performed using the Phoenix MICRON® IV retinal imaging microscope, and retinal thickness was quantified by manual segmentation. Blood pressure was measured using tail-cuff (BP-2010, Softron) after 7 days acclimation. Corticosterone was measured using QuicKey Pro Mouse CORT ELISA (Cat# E-OSEL-M0001) from morning serum. Complete blood counts were measured using Sysmex XN-10(B4).

### Statistical analysis

Unless stated otherwise, data are presented as mean ± s.d. or median (IQR). Normality was assessed using Shapiro–Wilk tests. Two-group comparisons used two-tailed Student’s t-tests (Welch correction when variances were unequal) or Mann–Whitney U tests. Multi-group comparisons used one- or two-way ANOVA with post hoc correction (Tukey or Bonferroni) or Kruskal–Wallis with Dunn correction. Longitudinal blood/urine Pb ratio was analyzed using linear mixed-effects models implemented in lme4^86^, with p-values from lmerTest^87^. The primary model specified fixed effects for time (days), genotype, and their interaction, and a random intercept for mouse ID: log2(ratio) ∼ time × genotype + (1 | mouse). Missing observations were handled by maximum likelihood (no imputation). Multiple-testing correction used Benjamini–Hochberg FDR where applicable (threshold 0.05). Exact *P* values are reported where relevant.

## Acknowledgments

We thank Professor Ying Shen and Professor Weiqiang Lin of the International Medical School, International Institutes of Medicine, The Fourth Affiliated Hospital, Zhejiang University School of Medicine, for their guidance on the brain- and kidney-related aspects of this project.We thank Yuchao Sun, Ying Liu, Lejun Zhang, Yuting Wang, Wenying Long and Yao Wu from the Core Facilities of the Fourth Affiliated Hospital, Zhejiang University School of Medicine and the International Institutes of Medicine, Zhejiang University, for technical assistance. We also thank Nijie, Beijia Wang, Yulan Zheng and Weiwei Jin from the Laboratory Animal Center of the International Institutes of Medicine, Zhejiang University, for meticulous animal care. We thank the Department of Laboratory Medicine and the Pathology Core of the Fourth Affiliated Hospital, Zhejiang University School of Medicine, and members of the Li laboratory for technical support and discussions. We thank Xiaoxia Wan from the Center of Cryo-Electron Microscopy (CCEM), Zhejiang University, for assistance with cryo-transmission electron microscopy. We thank GemPharmatech for assistance with mouse provision, IVF and microinjection procedures, Annoroad for kidney single-cell sequencing services, and Shanghai Chemlab Instrument Co., Ltd. for laser ablation instrumentation.

## Author Contributions

Chenyi Bo Zhang, Xiaoping Xia and Zhixiang Guo collected clinical samples and performed serum and urine biochemical analyses and complete blood counts. Chenyi Bo Zhang measured Pb content and conducted behavioral assays. Chenyi Bo Zhang, Xuliang Ma, Yukai Jason Zhong and Jingwen Luo performed mouse experiments. Jingwen Luo and Liang Xu Ma conducted intestinal permeability assays. Jiaoyan Ma performed mitochondrial function assays. Banghua He performed Western blot analyses. Huali Luo, Chenyi Bo Zhang and Haifeng Zhang conducted H&E and immunofluorescence staining. Qin Li performed genotyping of F-generation injection mice. Hongxiao Cui constructed small RNA libraries from kidney and testis samples. Guangnan Li constructed sperm RNA libraries. Chenyu Pei analyzed single-cell RNA sequencing data from testes and kidneys. Chen Zhang performed CRISPR-based whole-genome knockout screening in 293T cells with Pb treatment. Chenyi Bo Zhang and Xin Zhiguo Li analyzed the data. Xin Zhiguo Li conceived and designed the study. All authors contributed to manuscript preparation and approved the final version.

## Author Information

The authors declare no competing financial interests. Correspondence and requests for materials should be addressed to

## Data availability

Raw single-cell RNA sequencing data have been deposited in the China National Center for Bioinformation (CNCB) under accession number PRJCA058345. Processed gene expression matrices and associated metadata are provided as Extended Data files.

## Extended Data Materials

Extended Data Table 1-2

Extended Data Fig. 1-6

## References

1. Rout, P. & Jialal, I. in StatPearls (StatPearls Publishing, Treasure Island (FL), 2025).

2. Lefmann, T. & Combs-Orme, T. Prenatal stress, poverty, and child outcomes. Child and Adolescent Social Work Journal 31, 577–590 (2014).

3. Rees, N. & Fuller, R. The toxic truth: children’s exposure to lead pollution undermines a generation of future potential (Unicef, 2020).

4. Sen, A. et al. Multigenerational epigenetic inheritance in humans: DNA methylation changes associated with maternal exposure to lead can be transmitted to the grandchildren. Sci Rep 5, 14466 (2015).

5. Sobolewski, M. et al. Lineage- and Sex-Dependent Behavioral and Biochemical Transgenerational Consequences of Developmental Exposure to Lead, Prenatal Stress, and Combined Lead and Prenatal Stress in Mice. Environ Health Perspect 128, 27001 (2020).

6. Adokwe, J. B. et al. Concurrent Lead and Cadmium Exposure Among Diabetics: A Case-Control Study of Socio-Demographic and Consumption Behaviors. Nutrients 17, (2025).

7. Numan, A. T., Jawad, N. K. & Fawzi, H. A. Biochemical study of the effect of lead exposure in nonobese gasoline station workers and risk of hyperglycemia: A retrospective case-control study. Medicine (Baltimore) 103, e39152 (2024).

8. Jin, T., Park, E. Y., Kim, B. & Oh, J. K. Environmental exposure to lead and cadmium are associated with triglyceride glucose index. Sci Rep 14, 2496 (2024).

9. Yu, B. et al. Lead exposure and physical frailty in patients with type 2 diabetes mellitus: cross-sectional results from the METAL study. Endocrine 87, 987–996 (2025).

10. Lin, J. L. et al. Environmental exposure to lead and progressive diabetic nephropathy in patients with type II diabetes. Kidney Int 69, 2049–2056 (2006).

11. Huang, W. H. et al. Environmental lead exposure accelerates progressive diabetic nephropathy in type II diabetic patients. Biomed Res Int 2013, 742545 (2013).

12. Yamazaki, M. et al. Segmentation of the Pathophysiological Stages of Diabetic Changes in the db/db Mouse. J Toxicol Pathol 22, 133–137 (2009).

13. Cory-Slechta, D. A., Weiss, B. & Cox, C. Performance and exposure indices of rats exposed to low concentrations of lead. Toxicol Appl Pharmacol 78, 291–299 (1985).

14. Sobolewski, M. et al. Developmental Lead Exposure and Prenatal Stress Result in Sex-Specific Reprograming of Adult Stress Physiology and Epigenetic Profiles in Brain. Toxicol Sci 163, 478–489 (2018).

15. Flora, G., Gupta, D. & Tiwari, A. Toxicity of lead: A review with recent updates. Interdiscip Toxicol 5, 47–58 (2012).

16. Sanders, T., Liu, Y., Buchner, V. & Tchounwou, P. B. Neurotoxic effects and biomarkers of lead exposure: a review. Rev Environ Health 24, 15–45 (2009).

17. Toxicological Profile for Lead. (2020).

18. Gapp, K. et al. Implication of sperm RNAs in transgenerational inheritance of the effects of early trauma in mice. Nat Neurosci 17, 667–669 (2014).

19. Rodgers, A. B., Morgan, C. P., Leu, N. A. & Bale, T. L. Transgenerational epigenetic programming via sperm microRNA recapitulates effects of paternal stress. Proc Natl Acad Sci U S A 112, 13699–13704 (2015).

20. Chen, Q. et al. Sperm tsRNAs contribute to intergenerational inheritance of an acquired metabolic disorder. Science 351, 397–400 (2016).

21. Sharma, U. et al. Biogenesis and function of tRNA fragments during sperm maturation and fertilization in mammals. Science 351, 391–396 (2016).

22. Silbergeld, E. K. et al. Lead in bone: storage site, exposure source, and target organ. Neurotoxicology 14, 225–236 (1993).

23. Ingalls, A. M., Dickie, M. M. & Snell, G. D. Obese, a new mutation in the house mouse. J Hered 41, 317–318 (1950).

24. Zhang, Y. et al. Positional cloning of the mouse obese gene and its human homologue. Nature 372, 425–432 (1994).

25. Rader, J. I., Peeler, J. T. & Mahaffey, K. R. Comparative toxicity and tissue distribution of lead acetate in weanling and adult rats. Environ Health Perspect 42, 187–195 (1981).

26. Yamamoto, T. et al. Disturbed gastrointestinal motility and decreased interstitial cells of Cajal in diabetic db/db mice. J Gastroenterol Hepatol 23, 660–667 (2008).

27. Furuse, M., Sasaki, H., Fujimoto, K. & Tsukita, S. A single gene product, claudin-1 or -2, reconstitutes tight junction strands and recruits occludin in fibroblasts. J Cell Biol 143, 391–401 (1998).

28. Günzel, D. & Yu, A. S. Claudins and the modulation of tight junction permeability. Physiol Rev 93, 525–569 (2013).

29. Brun, P. et al. Increased intestinal permeability in obese mice: new evidence in the pathogenesis of nonalcoholic steatohepatitis. Am J Physiol Gastrointest Liver Physiol 292, G518–25 (2007).

30. Lee, B., Moon, K. M. & Kim, C. Y. Tight Junction in the Intestinal Epithelium: Its Association with Diseases and Regulation by Phytochemicals. J Immunol Res 2018, 2645465 (2018).

31. Young, C. F., Moussa, M. & Shubrook, J. H. Diabetic Gastroparesis: A Review. Diabetes Spectr 33, 290–297 (2020).

32. Yuan, J. H. et al. Impaired intestinal barrier function in type 2 diabetic patients measured by serum LPS, Zonulin, and IFABP. J Diabetes Complications 35, 107766 (2021).

33. Goyer, R. A. Mechanisms of lead and cadmium nephrotoxicity. Toxicol Lett 46, 153–162 (1989).

34. Sabath, E. & Robles-Osorio, M. L. Renal health and the environment: heavy metal nephrotoxicity. Nefrologia 32, 279–286 (2012).

35. Levey, A. S. et al. Definition and classification of chronic kidney disease: a position statement from Kidney Disease: Improving Global Outcomes (KDIGO). Kidney Int 67, 2089–2100 (2005).

36. Sembach, F. E. et al. Rodent models of diabetic kidney disease: human translatability and preclinical validity. Drug Discov Today 26, 200–217 (2021).

37. Aposhian, H. V. DMSA and DMPS--water soluble antidotes for heavy metal poisoning. Annu Rev Pharmacol Toxicol 23, 193–215 (1983).

38. Khalil-Manesh, F., Gonick, H. C., Cohen, A., Bergamaschi, E. & Mutti, A. Experimental model of lead nephropathy. II. Effect of removal from lead exposure and chelation treatment with dimercaptosuccinic acid (DMSA). Environ Res 58, 35–54 (1992).

39. Casagrande, S. S. & Gary-Webb, T. L. in Diabetes in America (eds Lawrence, J. M., Casagrande, S. S., Herman, W. H., Wexler, D. J. & Cefalu, W. T.) (National Institute of Diabetes and Digestive and Kidney Diseases (NIDDK), Bethesda (MD), 2023).

40. Cunningham, M. Chronic occupational lead exposure: the potential effect on sexual function and reproductive ability in male workers. AAOHN J 34, 277–279 (1986).

41. Lancranjan, I., Popescu, H. I., GAvănescu, O., Klepsch, I. & Serbănescu, M. Reproductive ability of workmen occupationally exposed to lead. Arch Environ Health 30, 396–401 (1975).

42. Corpas, I. et al. Testicular alterations in rats due to gestational and early lactational administration of lead. Reprod Toxicol 9, 307–313 (1995).

43. Patrick, L. Lead toxicity, a review of the literature. Part 1: Exposure, evaluation, and treatment. Altern Med Rev 11, 2–22 (2006).

44. Skinner, M. K., Manikkam, M. & Guerrero-Bosagna, C. Epigenetic transgenerational actions of environmental factors in disease etiology. Trends Endocrinol Metab 21, 214–222 (2010).

45. Heard, E. & Martienssen, R. A. Transgenerational epigenetic inheritance: myths and mechanisms. Cell 157, 95–109 (2014).

46. Nilsson, E. E. & Skinner, M. K. Environmentally Induced Epigenetic Transgenerational Inheritance of Reproductive Disease. Biol Reprod 93, 145 (2015).

47. Thomas, M. C., et al. Diabetic kidney disease. Nat Rev Dis Primers 1, 15018 (2015).

48. Tuttle, K. R. et al. Diabetic kidney disease: a report from an ADA Consensus Conference. Diabetes Care 37, 2864–2883 (2014).

49. Alpers, C. E. & Hudkins, K. L. Mouse models of diabetic nephropathy. Curr Opin Nephrol Hypertens 20, 278–284 (2011).

50. Chaudhuri, A., Ghanim, H. & Arora, P. Improving the residual risk of renal and cardiovascular outcomes in diabetic kidney disease: A review of pathophysiology, mechanisms, and evidence from recent trials. Diabetes Obes Metab 24, 365–376 (2022).

51. Cate, M. Health System Resource Allocation Models for SGLT2 Inhibitors in Low-and Middle-Income Countries. (2025).

52. Ben Maamar, M., Nilsson, E., Thorson, J. L. M., Beck, D. & Skinner, M. K. Transgenerational disease specific epigenetic sperm biomarkers after ancestral exposure to dioxin. Environ Res 192, 110279 (2021).

53. Nilsson, E. et al. Vinclozolin induced epigenetic transgenerational inheritance of pathologies and sperm epimutation biomarkers for specific diseases. PLoS One 13, e0202662 (2018).

54. Zatecka, E. et al. The Transgenerational Transmission of the Paternal Type 2 Diabetes-Induced Subfertility Phenotype. Front Endocrinol (Lausanne) 12, 763863 (2021).

55. Luo, J., Zhang, Y. & Luo, Z. Assessing the global burden of Type 2 diabetes in women of reproductive age. PLoS One 20, e0322787 (2025).

56. American, D. A. P. P. C. 2. Diagnosis and Classification of Diabetes: Standards of Care in Diabetes-2025. Diabetes Care 48, S27–S49 (2025).

57. Inker, L. A. et al. New Creatinine- and Cystatin C-Based Equations to Estimate GFR without Race. N Engl J Med 385, 1737–1749 (2021).

58. Saleem, M. et al. Heavy Metal(oid)s Contamination and Potential Ecological Risk Assessment in Agricultural Soils. J Xenobiot 14, 634–650 (2024).

59. Benedé-Ubieto, R., Estévez-Vázquez, O., Ramadori, P., Cubero, F. J. & Nevzorova, Y. A. Guidelines and Considerations for Metabolic Tolerance Tests in Mice. Diabetes Metab Syndr Obes 13, 439–450 (2020).

60. Chen, B. et al. Maternal inheritance of glucose intolerance via oocyte TET3 insufficiency. Nature 605, 761–766 (2022).

61. Amira, S., Soufane, S. & Gharzouli, K. Effect of sodium fluoride on gastric emptying and intestinal transit in mice. Experimental and Toxicologic Pathology 57, 59–64 (2005).

62. Thaiss, C. A. et al. Hyperglycemia drives intestinal barrier dysfunction and risk for enteric infection. Science 359, 1376–1383 (2018).

63. Scarfe, L. et al. Transdermal Measurement of Glomerular Filtration Rate in Mice. J Vis Exp (2018).

64. Schock-Kusch, D. et al. Transcutaneous measurement of glomerular filtration rate using FITC-sinistrin in rats. Nephrol Dial Transplant 24, 2997–3001 (2009).

65. Kyrylkova, K., Kyryachenko, S., Leid, M. & Kioussi, C. Detection of apoptosis by TUNEL assay. Methods Mol Biol 887, 41–47 (2012).

66. Schlatt, S. & Weinbauer, G. F. Immunohistochemical localization of proliferating cell nuclear antigen as a tool to study cell proliferation in rodent and primate testes. Int J Androl 17, 214–222 (1994).

67. Zheng, G. X. et al. Massively parallel digital transcriptional profiling of single cells. Nat Commun 8, 14049 (2017).

68. Fleming, S. J. et al. Unsupervised removal of systematic background noise from droplet-based single-cell experiments using CellBender. Nat Methods 20, 1323–1335 (2023).

69. Wolf, F. A., Angerer, P. & Theis, F. J. SCANPY: large-scale single-cell gene expression data analysis. Genome Biol 19, 15 (2018).

70. Germain, P. L., Lun, A., Garcia Meixide, C., Macnair, W. & Robinson, M. D. Doublet identification in single-cell sequencing data using scDblFinder. F1000Res 10, 979 (2021).

71. Tirosh, I. et al. Dissecting the multicellular ecosystem of metastatic melanoma by single-cell RNA-seq. Science 352, 189–196 (2016).

72. Stuart, T. et al. Comprehensive Integration of Single-Cell Data. Cell 177, 1888–1902.e21 (2019).

73. Korsunsky, I. et al. Fast, sensitive and accurate integration of single-cell data with Harmony. Nat Methods 16, 1289–1296 (2019).

74. McInnes, L., Healy, J. & Melville, J. Umap: Uniform manifold approximation and projection for dimension reduction. *arXiv preprint arXiv:1802.03426* (2018).

75. Traag, V. A., Waltman, L. & van Eck, N. J. From Louvain to Leiden: guaranteeing well-connected communities. Sci Rep 9, 5233 (2019).

76. Lotfollahi, M. et al. Mapping single-cell data to reference atlases by transfer learning. Nat Biotechnol 40, 121–130 (2022).

77. Xu, C. et al. Probabilistic harmonization and annotation of single-cell transcriptomics data with deep generative models. Mol Syst Biol 17, e9620 (2021).

78. Green, C. D. et al. A Comprehensive Roadmap of Murine Spermatogenesis Defined by Single-Cell RNA-Seq. Dev Cell 46, 651–667.e10 (2018).

79. Aibar, S. et al. SCENIC: single-cell regulatory network inference and clustering. Nat Methods 14, 1083–1086 (2017).

80. Finak, G. et al. MAST: a flexible statistical framework for assessing transcriptional changes and characterizing heterogeneity in single-cell RNA sequencing data. Genome Biol 16, 278 (2015).

81. Sun, Y. H. et al. Single-molecule long-read sequencing reveals a conserved intact long RNA profile in sperm. Nat Commun 12, 1361 (2021).

82. Seibenhener, M. L. & Wooten, M. C. Use of the Open Field Maze to measure locomotor and anxiety-like behavior in mice. J Vis Exp e52434 (2015).

83. Walf, A. A. & Frye, C. A. The use of the elevated plus maze as an assay of anxiety-related behavior in rodents. Nat Protoc 2, 322–328 (2007).

84. Leger, M. et al. Object recognition test in mice. Nat Protoc 8, 2531–2537 (2013).

85. Cruz-Sanchez, A. et al. Developmental onset distinguishes three types of spontaneous recognition memory in mice. Sci Rep 10, 10612 (2020).

86. Bates, D., Mächler, M., Bolker, B. & Walker, S. Fitting linear mixed-effects models using lme4. Journal of statistical software 67, 1–48 (2015).

87. Kuznetsova, A., Brockhoff, P. B. & Christensen, R. H. B. lmerTest package: tests in linear mixed effects models. Journal of statistical software 82, 1–26 (2017).

