## Extended data for "Type 2 diabetes amplifies environmental Pb toxicity and intergenerational risk"

**This PDF file includes:**

Caption for Extended Data Table 1-2
Extended Data Fig. 1-6

**Other Supplemental Materials for this manuscript includes the following:**

Extended Data Table 1 (separate file)
Extended Data Table 2 (separate file)

Detailed information and statistics for the sequencing data used in this study.

**Extended Data Table 1 (separate file)**

**Multi-element profiling and environmental Pb measurements.**

(A) Whole-blood concentrations of 27 elements in individuals with type 2 diabetes (T2D) and age- and sex-matched non-diabetic controls, determined by inductively coupled plasma–mass spectrometry (ICP–MS).

(B) Pb concentrations in drinking water collected from three independent municipal reservoirs supplying the Yiwu study region.

(C) Pb concentrations measured in ten sugar-free beverages (three independent batches per product).

(D) Pb concentrations measured in lip products from ten leading brands.

**Extended Data Table 2 (separate file)**

**Single-cell RNA sequencing quality control and metadata.**

(A) Number of cells and genes retained at each quality-control step for kidney single-cell RNA sequencing analysis in $F_0$, $F_1$ and $F_2$ mice.

(B) Metadata for the final analyzed kidney cell dataset across generations.

(C) Number of cells and genes retained at each quality-control step for testis single-cell RNA sequencing analysis in $F_0$ mice.

(D) Metadata for the final analyzed testis cell dataset.

**EXTENDED DATA FIGURE LEGENDS**

Zhang et al, Extended Data Fig. 1

**a**

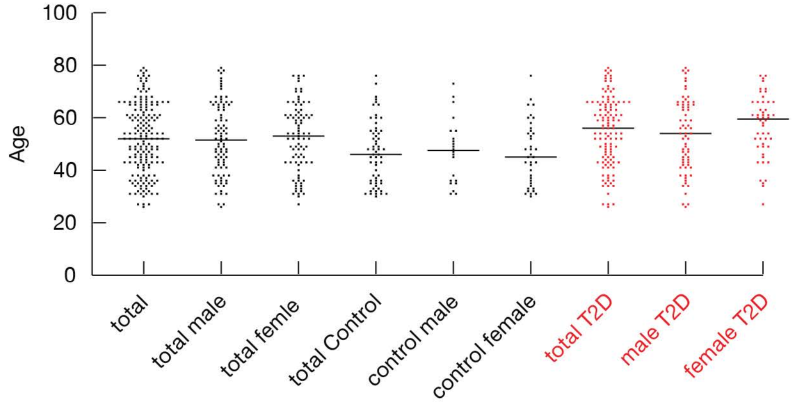


**b**

Spearman correlation analysis in T2D populations

| | Age | BMI | Hemo-globin A1 | Fasting blood glucose | Triglycerides | Cholesterol | HDL-C | LDL-C | Total bilirubin | Direct bilirubin | Indirect bilirubin | Total protein | Albumin | Globulin | Albumin-globulin ratio | ALT | AST | Alkaline phosphatase | Urea | Creatinine | Uric acid | estiated glomerular filtration rate(eGFR) |
|---|---|---|---|---|---|---|---|---|---|---|---|---|---|---|---|---|---|---|---|---|---|---|
| Correlation coefficient | 0.15 | 0.01 | 0.13 | -0.03 | 0.10 | 0.02 | 0.00 | -0.13 | -0.03 | -0.08 | 0.00 | 0.12 | -0.03 | 0.16 | -0.17 | -0.04 | 0.03 | 0.03 | 0.17 | .36** | .21* | -.34** |
| Sig.(two-tailed) | 0.11 | 0.91 | 0.21 | 0.78 | 0.33 | 0.81 | 1.00 | 0.20 | 0.76 | 0.46 | 0.99 | 0.23 | 0.77 | 0.12 | 0.10 | 0.69 | 0.75 | 0.76 | 0.10 | 0 | 0.04 | 0.00 |

* At the 0.05 level (two-tailed), the correlation is significant
** At the 0.01 level (two-tailed), the correlation is significant

Spearman correlation analysis in control populations

| | Age | BMI | Hemo-globin A1 | Fasting blood glucose | Triglycerides | Cholesterol | HDL-C | LDL-C | Total bilirubin | Direct bilirubin | Indirect bilirubin | Total protein | Albumin | Globulin | Albumin-globulin ratio | ALT | AST | Alkaline phosphatase | Urea | Creatinine | Uric acid | estiated glomerular filtration rate(eGFR) |
|---|---|---|---|---|---|---|---|---|---|---|---|---|---|---|---|---|---|---|---|---|---|---|
| Correlation coeffient | 0.14 | 0.42 | -0.22 | 0.14 | 0.07 | 0.11 | -0.09 | 0.17 | 0.08 | 0.09 | 0.05 | 0.10 | -0.01 | 0.09 | -0.02 | -0.07 | -0.08 | 0.06 | -0.00 | 0.17 | 0.15 | -0.25 |
| Sig.(two-tailed) | 0.29 | 0.11 | 0.10 | 0.29 | 0.63 | 0.41 | 0.49 | 0.21 | 0.55 | 0.51 | 0.73 | 0.47 | 0.93 | 0.49 | 0.89 | 0.62 | 0.54 | 0.68 | 0.98 | 0.21 | 0.02 | 0.06 |

* At the 0.05 level (two-tailed), the correlation is significant
** At the 0.01 level (two-tailed), the correlation is significant

**c**

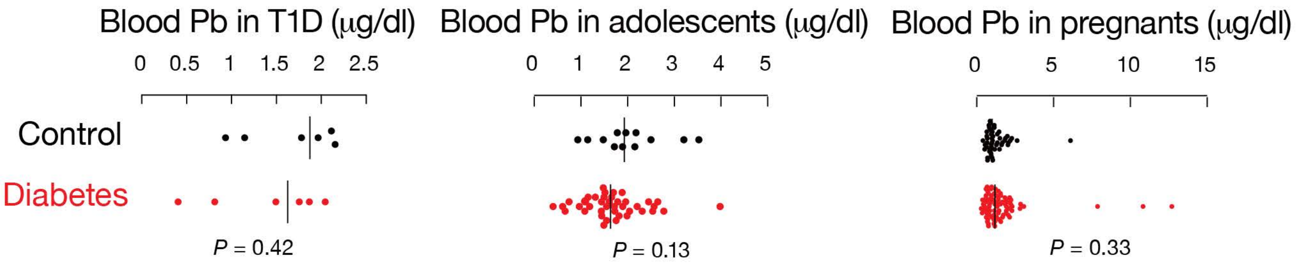


**d**

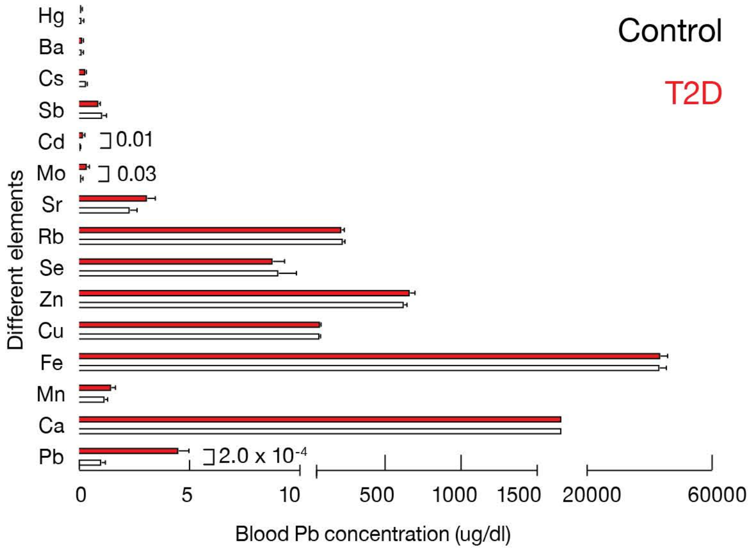

**Extended Data Fig. 1 | Human cohort characteristics and multi-element profiling.** **a**, Age distribution of all study participants, non-diabetic controls, and individuals with type 2 diabetes mellitus (T2D), shown for males and females separately. Data represent number of individuals per age bin (n = 61 controls; n = 118 T2D). **b**, Spearman rank correlation coefficients (ρ) between whole-blood Pb concentration and clinical parameters in non-diabetic controls and T2D individuals. Clinical parameters include age, body mass index (BMI), hemoglobin-A1(HbA1), fasting blood glucose, alanine aminotransferase (ALT), aspartate aminotransferase (AST), triglycerides (TG), total cholesterol (TC), serum creatinine (Cr), uric acid (UA), and estimated glomerular filtration rate (eGFR). Two-tailed Spearman correlation; exact *P* values indicated. **c**, Whole-blood Pb concentrations in additional human subgroups: type 1 diabetes mellitus (T1D), juvenile-onset T2D, and gestational diabetes mellitus (GDM), compared with age-matched non-diabetic controls (non-diabetic controls n=6, 12, 49 per group; diabetic group n = 6, 40, 79 per group). Two-tailed Mann–Whitney U test. **d**, Multi-element inductively coupled plasma–mass spectrometry (ICP–MS) profiling in a randomly selected subset of participants (non-diabetic controls n = 6; T2D n = 24). Of 27 screened elements, 15 were above the limit of quantification (LOQ) in ≥80% of samples and included in statistical comparison. Data are mean ± standard error of the mean (s.e.m.). Two-tailed Mann–Whitney U test.

Zhang et al, Extended Data Fig. 2

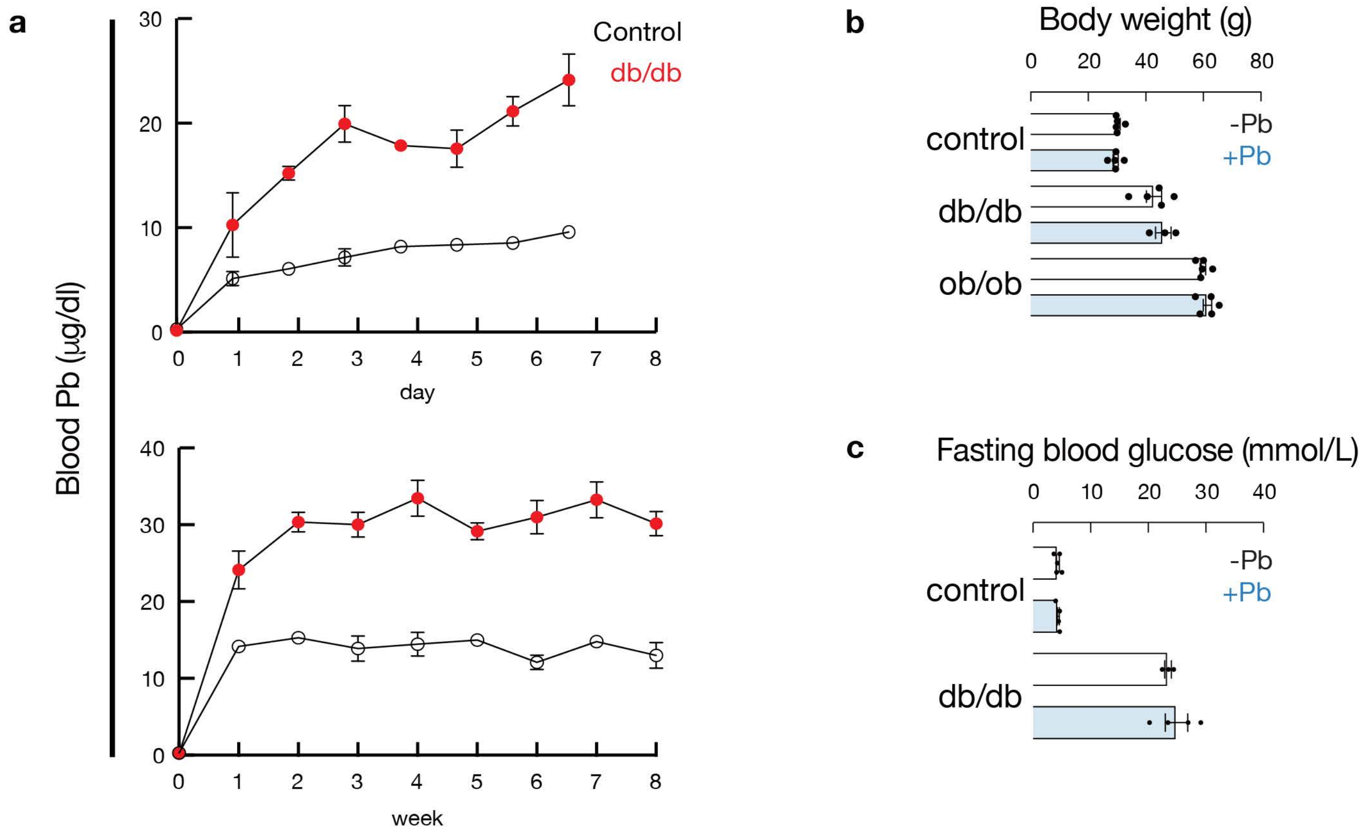

**Extended Data Fig. 2 | Pb kinetics and metabolic parameters during chronic exposure. a**, Whole-blood Pb concentration measured daily during days 1–7 and at week 8 of exposure to 100 parts per million (ppm) lead(II) acetate in drinking water in non-diabetic controls (m/m) and db/db mice (n = 3-4 per group). Data are mean ± s.e.m. **b**, Body weight after 8 weeks of Pb exposure in m/m, m/m(Pb), db/db, and db/db(Pb) mice (n = 3-5 per group). Two-way analysis of variance (ANOVA). **c**, Fasting blood glucose measured at week 16 following a 16-hour fast (n = 3-5 per group). Two-way ANOVA.

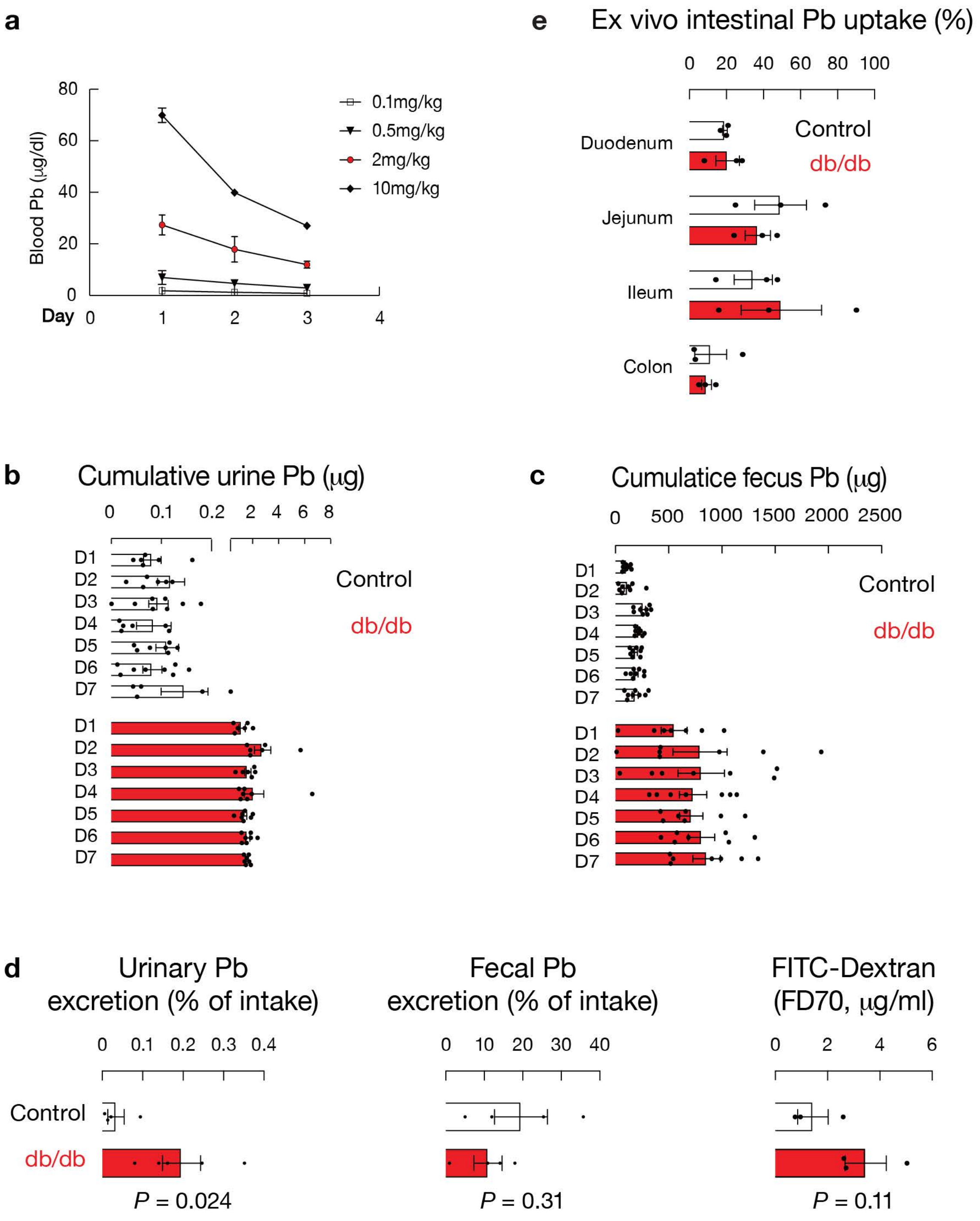
a
Blood Pb (μg/dl)
80
60
40
20
0
Day 0
1
2
3
4
0.1mg/kg
0.5mg/kg
2mg/kg
10mg/kg
e
Ex vivo intestinal Pb uptake (%)
0 20 40 60 80 100
Duodenum
Jejunum
Ileum
Colon
Control
db/db
b
Cumulative urine Pb (μg)
0 0.1 0.2 2 4 6 8
D1
D2
D3
D4
D5
D6
D7
Control
db/db
c
Cumulatice fecus Pb (μg)
0 500 1000 1500 2000 2500
D1
D2
D3
D4
D5
D6
D7
Control
db/db
d
Urinary Pb excretion (% of intake)
0 0.1 0.2 0.3 0.4
Control
db/db
P = 0.024
Fecal Pb excretion (% of intake)
0 10 20 30 40
P = 0.31
FITC-Dextran (FD70, μg/ml)
0 2 4 6
P = 0.11

**Extended Data Fig. 3 | Pb pharmacokinetics, excretion and intestinal barrier validation. a**, Whole-blood Pb concentration measured day1,2,3 after intravenous injection of Pb acetate at doses ranging from 0.5 to 10 mg $kg^{-1}$ in m/m mice (n = 3 per dose). Linear regression analysis shown. **b,c**, Cumulative urinary (b) and fecal (c) Pb excretion over 7 days of continuous exposure to 100 ppm lead(II) acetate in drinking water (n = 6-8 per group). **d**, fraction of Pb excreted in urine (left) and feces (middle) relative to ingested dose within 24 h after gavage; plasma fluorescence 30 min after gavage of 70 kDa FITC–dextran (FD70)(right). Data are mean ± s.e.m. Two-tailed unpaired t-test or two-way ANOVA as indicated. **e**, Ex vivo Pb uptake by isolated intestinal segments (duodenum, jejunum, ileum, colon) after 30 min luminal exposure (n = 3 per group). Data are mean ± s.e.m.

Zhang et al, Extended Data Fig. 4

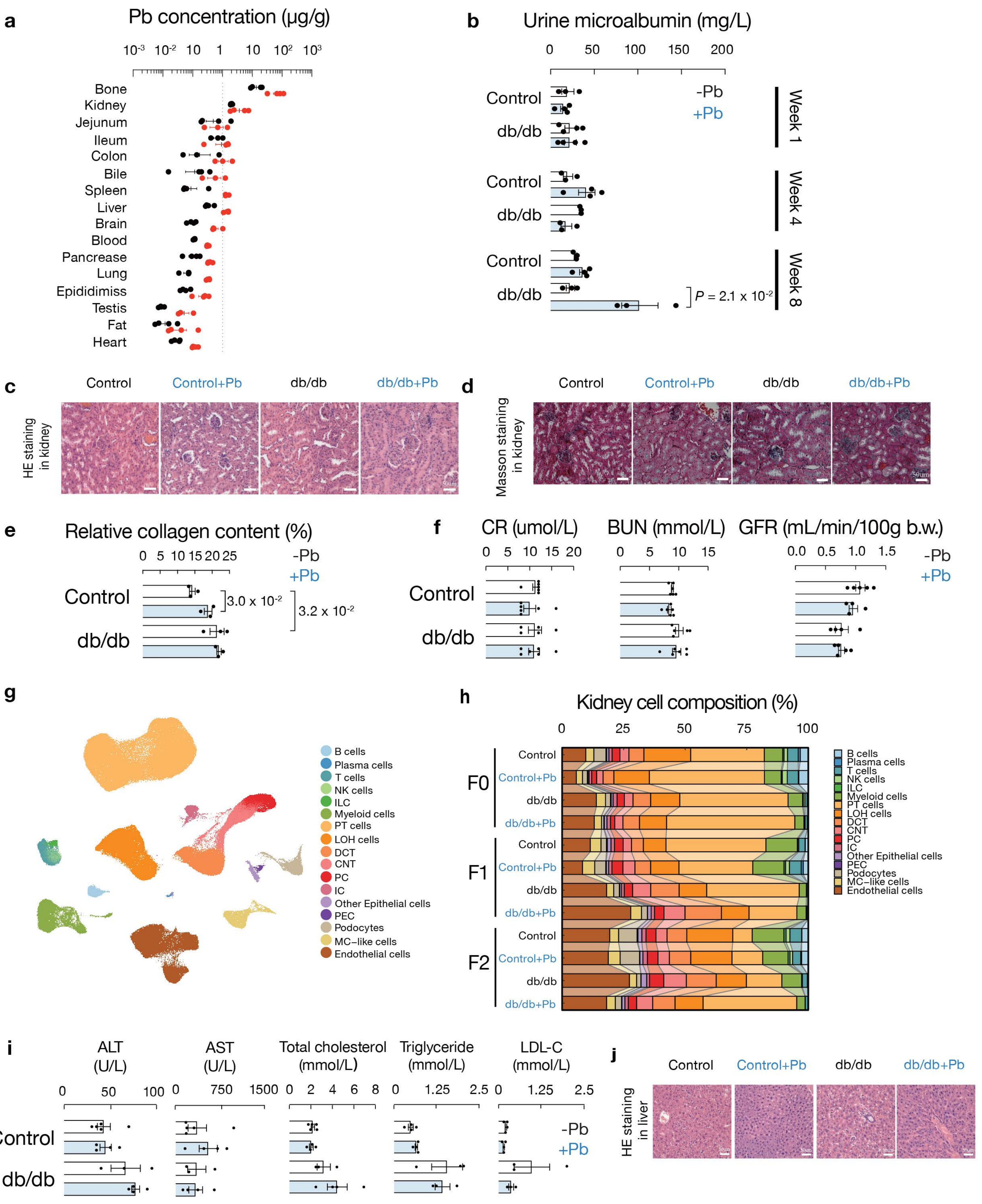

Zhang et al, Extended Data Fig. 4

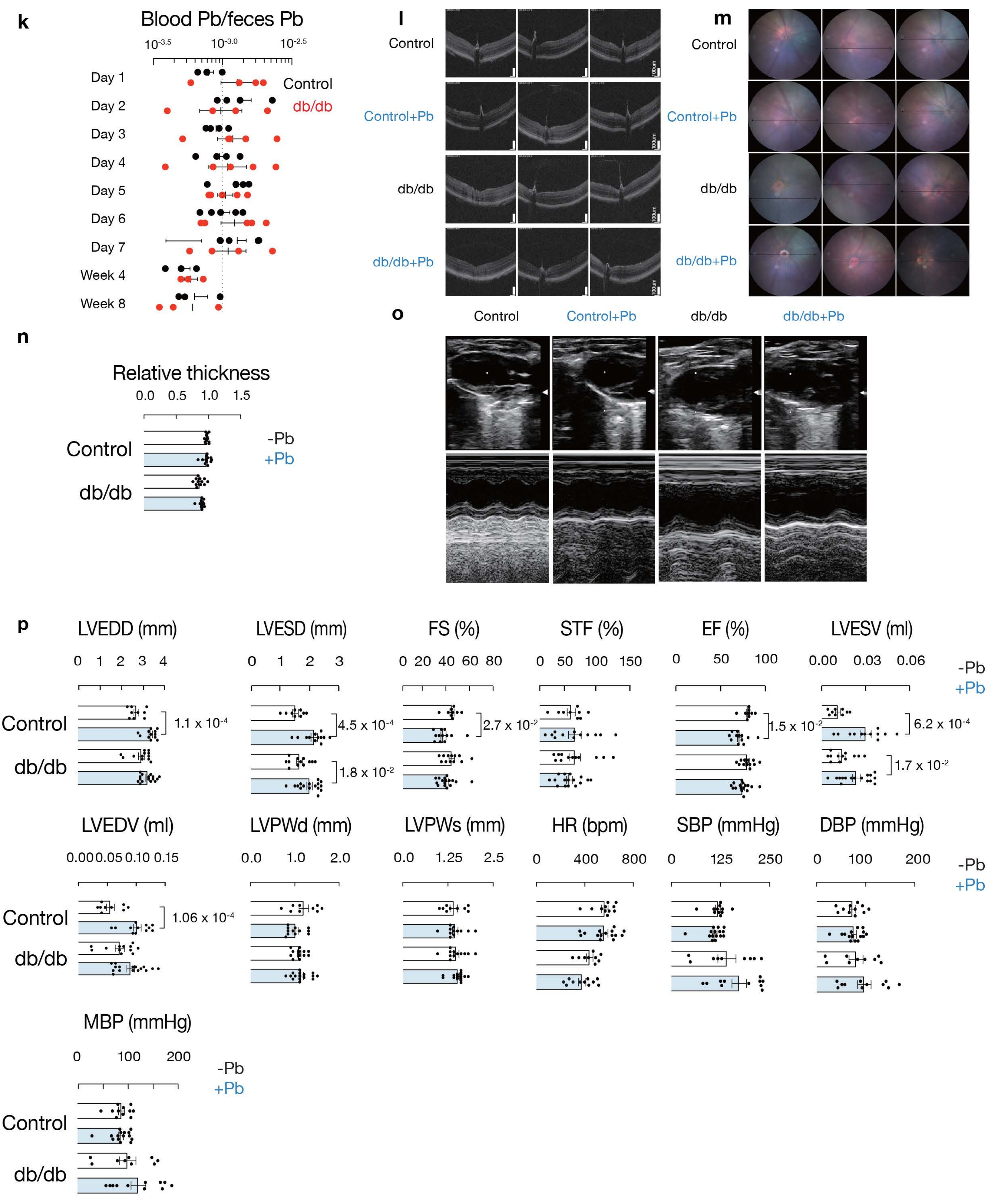

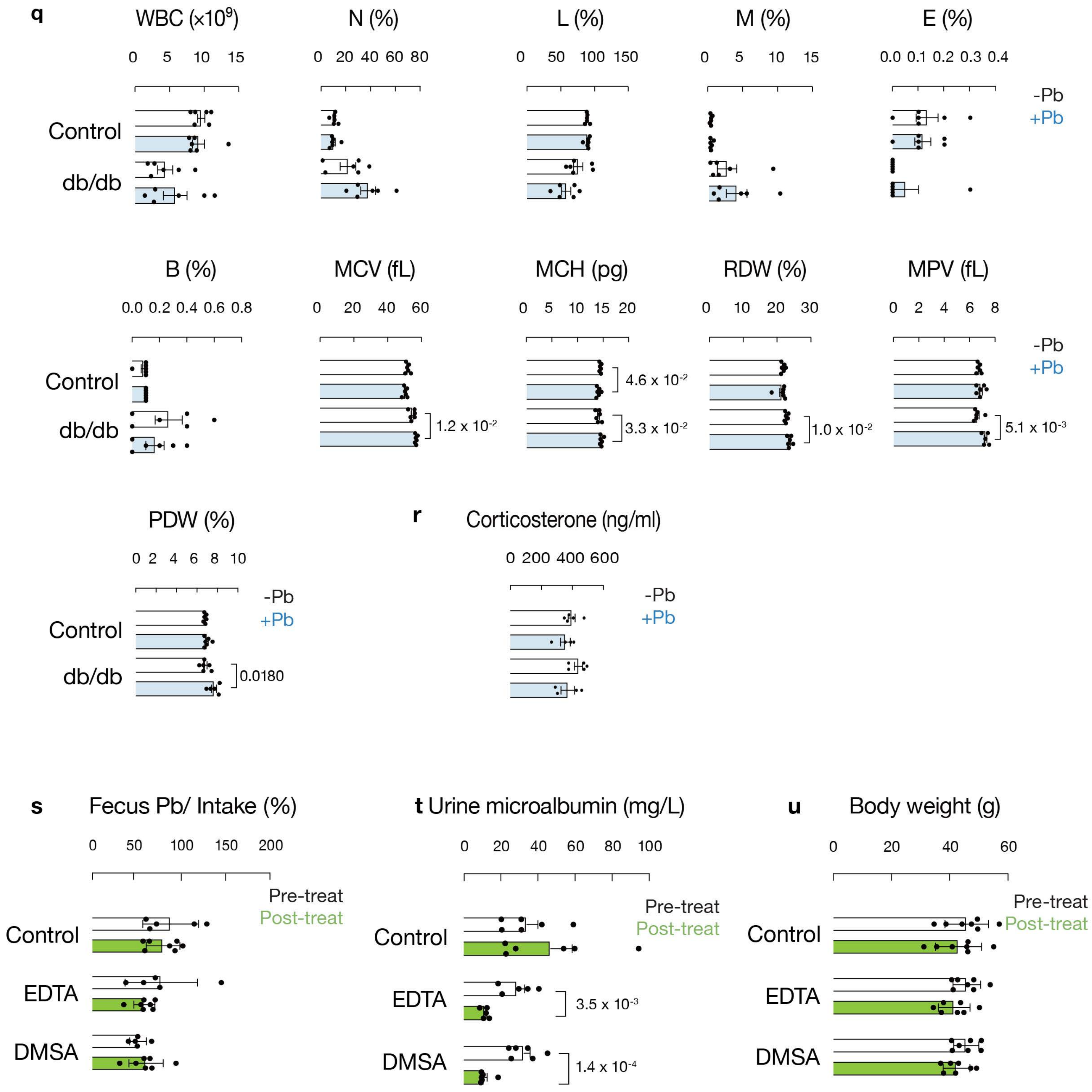
q
WBC (×10^9)
0 5 10 15
N (%)
0 20 40 60 80
L (%)
0 50 100 150
M (%)
0 5 10 15
E (%)
0.0 0.1 0.2 0.3 0.4
-Pb
+Pb
Control
db/db
B (%)
0.0 0.2 0.4 0.6 0.8
MCV (fL)
0 20 40 60
1.2 x 10-2
MCH (pg)
0 5 10 15 20
4.6 x 10-2
3.3 x 10-2
RDW (%)
0 10 20 30
1.0 x 10-2
MPV (fL)
0 2 4 6 8
5.1 x 10-3
-Pb
+Pb
Control
db/db
PDW (%)
0 2 4 6 8 10
-Pb
+Pb
Control
db/db
0.0180
r
Corticosterone (ng/ml)
0 200 400 600
-Pb
+Pb
s
Fecus Pb/ Intake (%)
0 50 100 150 200
Pre-treat
Post-treat
Control
EDTA
DMSA
t Urine microalbumin (mg/L)
0 20 40 60 80 100
Pre-treat
Post-treat
Control
EDTA
3.5 x 10-3
DMSA
1.4 x 10-4
u
Body weight (g)
0 20 40 60
Pre-treat
Post-treat
Control
EDTA
DMSA

**Extended Data Fig. 4 | Organ Pb distribution, renal histology and systemic phenotyping.** **a**, Pb concentrations measured by ICP–MS in blood, bone, kidney, liver, and brain after 8 weeks of Pb exposure (n = 3-4 per group). Units: μg $g^{-1}$ tissue or μg $dL^{-1}$ blood. **b**, Urine microalbumin concentration (mg $L^{-1}$) at weeks 1, 4 and 8 (n = 3-4 per group). **c**, Representative hematoxylin and eosin (H&E)-stained kidney sections from m/m, m/m(Pb), db/db, and db/db(Pb) mice. Scale bar, 50 μm. **d**, Representative Masson's trichrome staining of kidney sections. **e**, Quantification of fibrotic area (percentage of cortical area) from Masson's trichrome staining (n = 3 per group). **f**, Serum creatinine (μmol $L^{-1}$), blood urea nitrogen (BUN; mmol $L^{-1}$), and estimated glomerular filtration rate (eGFR; mL $min^{-1}$ 1.73 $m^{-2}$) at week 8 (n = 4-6 per group). **g**, Uniform manifold approximation and projection (UMAP) visualization of single-cell RNA sequencing (scRNA-seq) data from kidney across m/m, m/m(Pb), db/db, and db/db(Pb) groups. **h**, Relative proportions of annotated kidney cell types across genotypes and generations. **i**, Serum ALT, AST, total cholesterol, triglycerides, and low-density lipoprotein (LDL) levels at week 8 (n = 3-4 per group). **j**, Representative H&E-stained liver sections. **k**, Ratio of blood Pb concentration to fecal Pb concentration over time (n = 3-5 per group). **l**, Spectral-domain optical coherence tomography (SD-OCT) images of retina at week 8. **m**, Fundus photographs of retina. **n**, Quantification of total retinal thickness (μm) (n = 12 per group). **o,p**, Echocardiographic parameters including left ventricular end-diastolic diameter (LVEDD), left ventricular end-systolic diameter (LVESD), fractional shortening (FS), systolic time fraction (STF), ejection fraction (EF), left ventricular end-systolic volume (LVESV),left ventricular end-diastolic volume (LVEDV), left ventricular posterior wall thickness in diastole (LVPWd), left ventricular posterior wall thickness in systole (LVPWs), heart rate(HR), systolic blood pressure(SBP), diastolic blood pressure(DBP), mean blood pressure(MBP) (n = 9 per group). **q**, Hematological parameters including white blood cell (WBC) count, neutrophil(N) percentage, lymphocyte(L) percentage, monocyte(M) percentage, eosinophil(E) percentage, basophil(B) percentage, mean corpuscular volume (MCV), mean corpuscular hemoglobin (MCH), red cell distribution width (RDW), mean platelet volume (MPV), and platelet distribution width (PDW) (n = 6 per group). **r**, Serum corticosterone measured from morning samples (9:00–11:00) (n = 4-5 per group). **s–u**, Chelation treatment outcomes: fraction of ingested Pb excreted in feces (**s**), urine microalbumin (**t**), and body weight (**u**) before and after 5-day treatment with EDTA or DMSA (n = 6-7 per group). Data are mean ± s.e.m. unless otherwise indicated.

Zhang et al, Extended Data Fig. 5

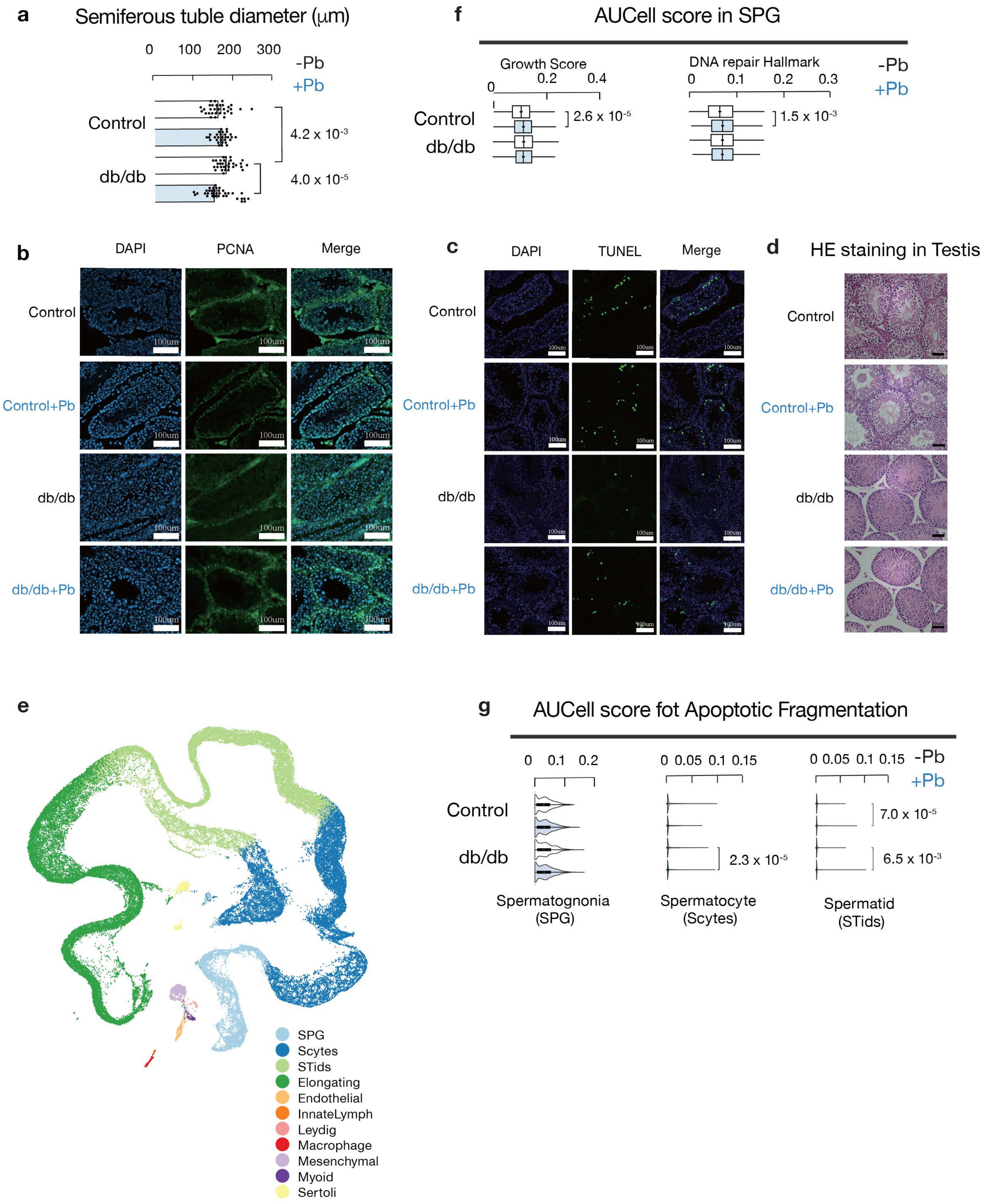

**Extended Data Fig. 5 | Testicular histology and single-cell transcriptomic validation.** **a**, Mean seminiferous tubule diameter (μm) measured from transverse sections (n = 30 per group). **b**, Representative proliferating cell nuclear antigen (PCNA) immunostaining of spermatogonia. **c**, Representative TUNEL (terminal deoxynucleotidyl transferase dUTP nick end labeling) staining of apoptotic germ cells. **d**, H&E-stained testis sections. **e**, UMAP visualization of testicular scRNA-seq data. **f**, Heat map of marker genes used for germ-cell and somatic-cell annotation. **g,** AUCell-based gene set activity scores for Apoptotic DNA Fragmentation (GO) in SPG, round spermatids (STids), and spermatocytes (Scytes). Statistical comparisons were performed using Wilcoxon test.Data are mean ± s.e.m. Two-way ANOVA or Mann–Whitney U test as appropriate.

Zhang et al, Extended Data Fig. 6

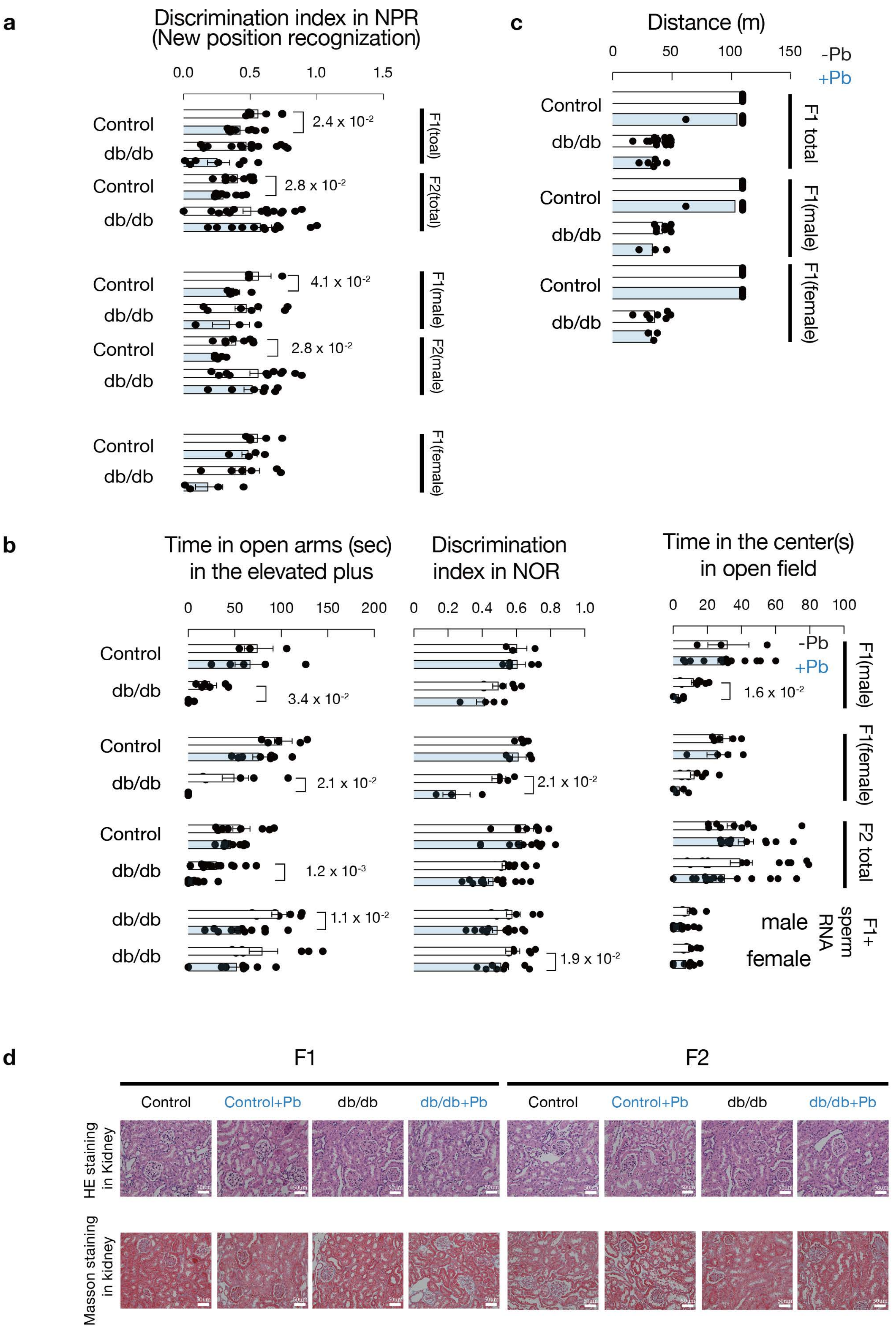

Zhang et al, Extended Data Fig. 6

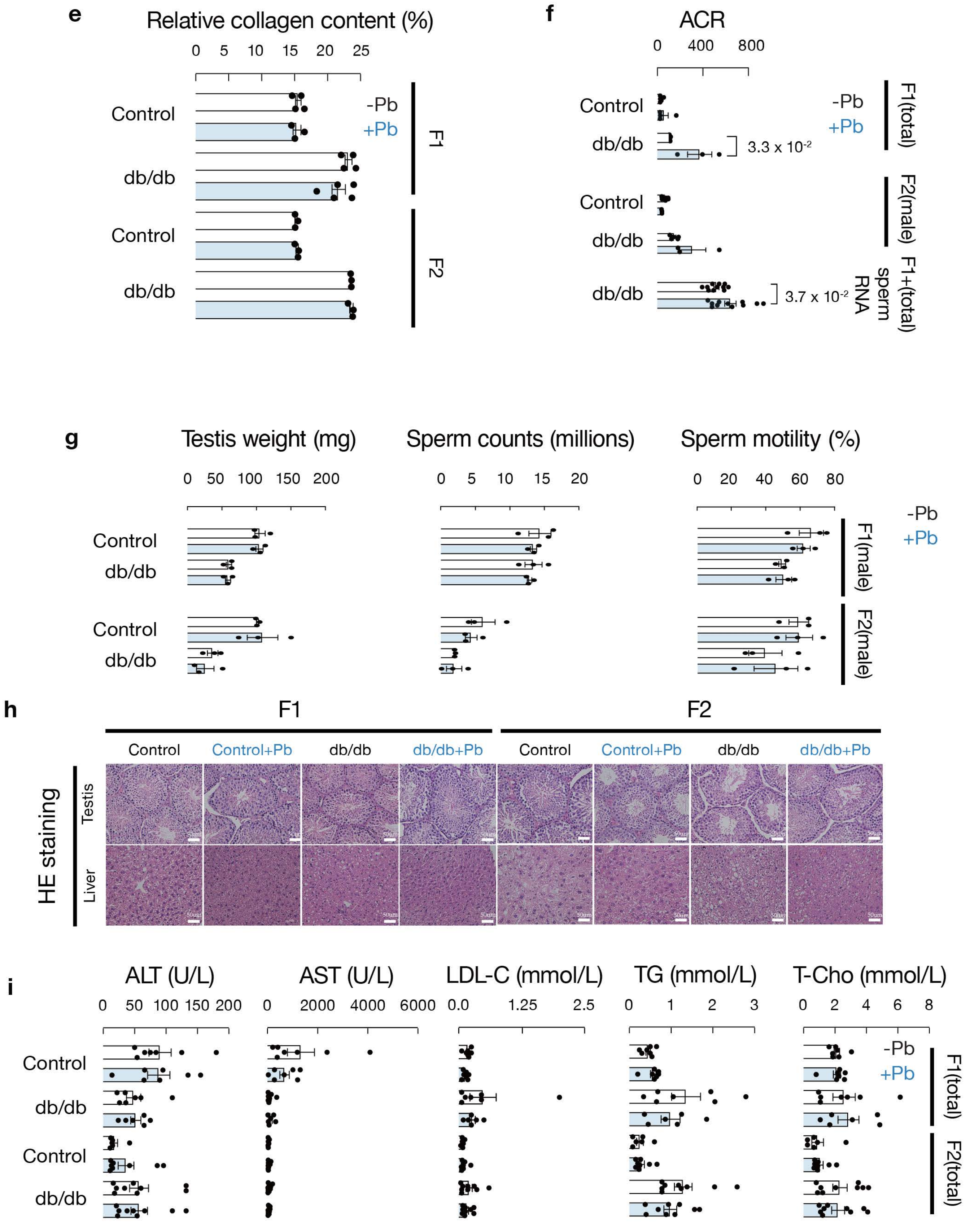

**Extended Data Fig. 6 | Behavioral and renal phenotypes in $F_1$ and $F_2$ offspring.** **a**, Novel position recognition test assessing spatial memory in $F_1$ and $F_2$ offspring (n = 8-18 per group). **b**, Behavioral testing (elevated plus maze, novel object recognition, open field) in $F_1$, $F_2$, and sperm RNA–injection offspring (n = 8-18 per group). **c**, Treadmill endurance performance in $F_1$ offspring (n = 6-15 per group). **d,** Kidney histology (upper) Masson's trichrome staining (lower) of kidney sections in F, F2 offspring. **e**, Quantification of fibrotic area (percentage of cortical area) from Masson's trichrome staining in $F_1$ and $F_2$ offspring (n = 3 per group). **f**, ACR in $F_1$(total), $F_2$(male), and sperm RNA–injection(total) offspring (n = 3-11per group). Data are mean ± s.e.m. Statistical tests as indicated. **g**, Testis weight, sperm count, and sperm motility in $F_1$ and F2 offspring (n = 3 per group). **h**, Representative H&E-stained testis sections from $F_1$ and F2 offspring. **i**, Serum ALT, AST, total cholesterol, triglycerides, and LDL in $F_1$ and F2 offspring.
